# Fine mapping without phenotyping: Identification of selection targets in secondary Evolve and Resequence experiments

**DOI:** 10.1101/2021.01.27.428395

**Authors:** Anna Maria Langmüller, Marlies Dolezal, Christian Schlötterer

## Abstract

Evolve and Resequence (E&R) studies investigate the genomic selection response of populations in an Experimental Evolution setup. Despite the popularity of E&R, empirical studies in sexually reproducing organisms typically suffer from an excess of candidate loci due to linkage disequilibrium, and single gene or SNP resolution is the exception rather than the rule. Recently, so-called “secondary E&R” has been suggested as promising experimental follow-up procedure to confirm putatively selected regions from a primary E&R study. Secondary E&R provides also the opportunity to increase mapping resolution by allowing for additional recombination events, which separate the selection target from neutral hitchhikers. Here, we use computer simulations to assess the effect of different crossing schemes, population size, experimental duration, and number of replicates on the power and resolution of secondary E&R. We find that the crossing scheme and population size are crucial factors determining power and resolution of secondary E&R: a simple crossing scheme with few founder lines consistently outcompetes crossing schemes where evolved populations from a primary E&R experiment are mixed with a complex ancestral founder population. Regardless of the experimental design tested, a population size of at least 4,800 individuals, which is roughly 5 times larger than population sizes in typical E&R studies, is required to achieve a power of at least 75%. Our study provides an important step towards improved experimental designs aiming to characterize causative SNPs in Experimental Evolution studies.

**Significance:** Despite the popularity of Evolve and Resequence (E&R) to investigate genomic selection responses, most studies that use sexually reproducing organisms have broad selection signatures and an excess of candidate loci due to linkage disequilibrium. In this study, we use computer simulations and statistical modelling to evaluate the effects of different experimental and population genetic parameters on the success of potential follow-up experiments (=secondary E&R) aiming to validate and fine-map selection signatures of primary studies. We found that a large population size in combination with a simple crossing scheme is key to the success of secondary E&R in *Drosophila*.

## Introduction

Deciphering the genetic architecture of adaptation is one of the longstanding goals in evolutionary biology. Experimental Evolution (EE) has become a popular approach to study adaptation in real time (Garland & Rose 2009; Kawecki et al. 2012). In contrast to natural populations, EE offers the key advantage of replicating experiments under controlled laboratory conditions (Schlötterer et al. 2015). Evolve and Resequence (E&R) (Turner et al. 2011; Long et al. 2015; Schlötterer et al. 2015) – a combination of EE with Next Generation Sequencing – facilitates in-depth analysis of the genomic responses to selection, with the ultimate goal to identify and characterize individual adaptive loci.

E&R has already been successful in investigating genomic selection responses from standing genetic variation in adapting sexually reproducing organisms, such as chicken (Johansson et al. 2010), yeast (Burke et al. 2014), and *Drosophila* (Teotónio et al. 2009; Remolina et al. 2012; Martins et al. 2014; Barghi et al. 2019). Despite its popularity, E&R typically suffers from an excess of candidates caused by linkage disequilibrium between true causative SNPs and neutral hitchhikers (Nuzhdin & Turner 2013; Tobler et al. 2014; Franssen et al. 2015), which decreases the resolution of E&R studies and makes single gene resolution (Martins et al. 2014) the exception rather than the rule.

The problem of candidate excess in E&R studies has been approached from different angles. More refined statistical tests have been developed (Topa et al. 2015; Iranmehr et al. 2017; Kelly & Hughes 2019; Spitzer et al. 2020), and the combination of time-series data with replicate populations has been identified as particularly powerful (Lang et al. 2013; Burke et al. 2014; Barghi et al. 2020). Organisms with a higher recombination rate and a lack of large segregating inversions that suppress recombination events have been suggested to be better suited for E&R studies (Barghi et al. 2017). Computer simulations showed that the power of E&R studies can be significantly improved by increasing the number of replicate populations, the experimental duration, or by adjusting the applied selection regime (Baldwin-Brown et al. 2014; Kofler & Schlötterer 2014; Kessner & Novembre 2015; Vlachos & Kofler 2019).

Burny et al. (2020) recently suggested an experimental follow-up procedure (“secondary E&R”) to validate selection signals of primary E&R studies. The basic idea of secondary E&R is that putative selection targets determined in the primary E&R study should rise in frequency again when exposed to the same environmental conditions during an additional E&R conducted after the primary experiment (Figure 1A). This experimental validation of selection signals is especially attractive before starting the time-consuming functional characterization of putatively selected alleles (e.g. based on the CRISPR/Cas technology) (Gratz et al. 2013). A hitherto under-explored potential of secondary E&R is that the additional recombination events during the secondary E&R can be used to fine map selection signals of primary experiments. In addition to mixing evolved genotypes of a primary E&R with non-adapted ancestral founder genotypes (coined “dilution” by Burny et al. (2020)), we propose several different secondary E&R crossing schemes for validating and fine-mapping of putative selection targets.

**Figure 1.**
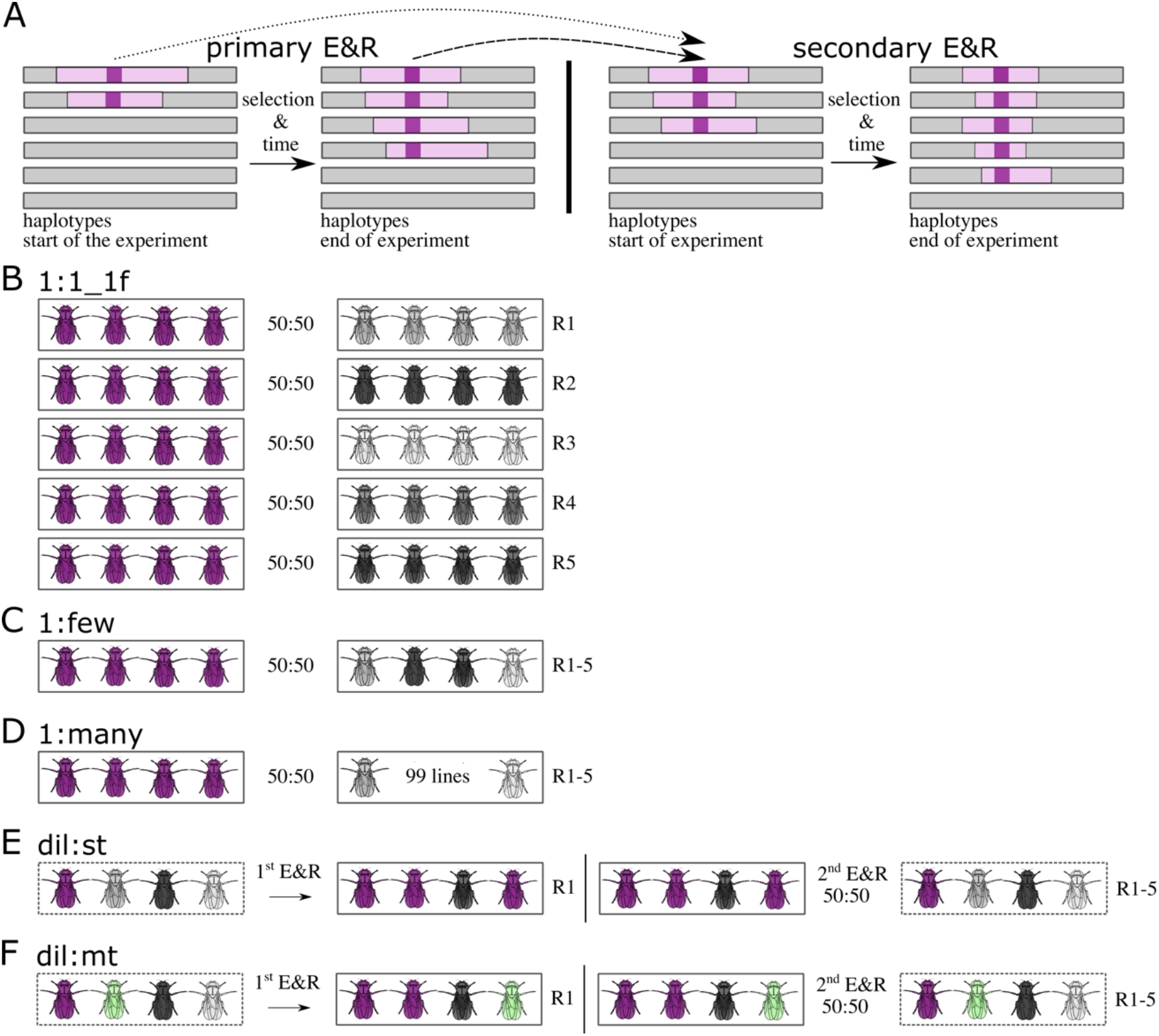
Basic idea of secondary Evolve and Resequence (E&R) and simulated crossing schemes. (A) The basic idea of secondary E&R is that a putative selection target (purple) determined in a primary E&R study (left) should rise in frequency again when exposed to the same environment (i.e., selection regime), during secondary E&R (right). Additional recombination events during the secondary E&R allow to fine map selected regions of primary E&R experiments (*i.e.,* reduce the number hitchhikers indicated in light pink) The two arrows indicate that the secondary E&R is either started with evolved haplotypes (dashed arrow) or with specific founder lines of the primary E&R (dotted arrow). (B) **1:1_1f** crossing scheme: Inbred flies with one target of selection (purple) are crossed to inbred flies without known beneficial variants. The starting frequency of each genotype is 50 %. In each replicate the line with the beneficial allele (focal line, purple) is crossed to a different line lacking beneficial mutations (non-focal lines, different shades of grey). (C) **1:few** crossing scheme: The focal line is crossed to a pool of flies with five different genotypes without known selection targets. The starting frequency of the focal line is again 50 %. (D) **1:many** crossing scheme: The focal line is crossed to a pool of 99 lines without known selection targets. (E) **dil:st** crossing scheme: 50 % of an evolved population originating from a primary E&R is replaced by ancestral genotypes of the primary E&R. The entire population has only one single target of selection (focal SNP, purple flies) (F) **dil:mt** crossing scheme: 50 % of an evolved population originating from a primary E&R is replaced by ancestral genotypes of the primary E&R. The ancestral population carries 16 targets of selection (flies carrying different beneficial SNPs are shown in purple, and green).

We evaluate the power and resolution of different secondary E&R designs to identify causative SNPs via extensive computer simulations. We use logistic regression to assess which simulated experimental and population genetic parameters have a significant effect on the success of secondary E&R. Selection coefficient, dominance coefficient, and mean starting allele frequency of the selection target all have a significant effect on the success of secondary E&R. However, crossing scheme and population size emerge as the most influential parameters. We show that the population size of secondary E&R experiments needs to be at least 5 times larger than currently used population sizes in typical primary E&R studies with *Drosophila* to achieve a power above 75%. Furthermore, we show that the crossing scheme is a crucial experimental parameter shaping the power of secondary E&R - a simple crossing scheme with few founder lines results in higher power and resolution compared to more complex crossing schemes.

## Material & Methods

### Outline of the simulation framework

The non-adapted ancestral founder genotypes used in our simulation study are a randomly chosen subset of 100 haplotypes from a panel of 189 sequenced *D. simulans* haplotypes originally collected in Tallahassee (Florida, USA) capturing the amount of standing genetic variation in a natural *Drosophila* population (Howie et al. 2019; Barghi et al. 2019). We use the term “founder line” for an inbred isofemale line homozygous for one of these ancestral haplotypes. In order to speed up the calculations, we only simulated chromosome-arm 2L. Simulations with linkage were conducted with MimicrEE2 (v206) (Vlachos & Kofler 2018) using the *D. simulans* recombination map (Howie et al. 2019). MimicrEE2 is a forward-simulation framework for E&R studies that can simulate evolving experimental populations based on their haplotype information and genome-wide recombination rates. We used the w-mode of MimicrEE2 which computes the fitness of individuals directly from the selection coefficients. If not stated otherwise, we simulated positive, additive selection for biallelic SNPs, with selection coefficients being uniformly sampled between 0.07 and 0.1 for each positively selected SNP and tracked the frequency of all SNPs over time (ranging from 479,507 to 970,466 SNPs, Table S1). We chose to simulate rather strong selection reasoning that alleles with a high selection coefficient are more likely to be experimentally tested. On the other hand, strongly selected alleles result in many neutral linked hitchhikers producing false positive signals that adversely impact the mapping resolution (Kofler & Schlötterer 2014) - which requires follow-up studies to identify the target of selection. If not stated otherwise, selected SNPs were randomly chosen with an equal probability to be either co-dominant (dominance coefficient h=0.5), or dominant (h=1). We did not consider recessive loci (h=0), because we do not anticipate that fully recessive targets would result in sufficiently large allele frequency changes to be detected in primary E&R experiments (Baldwin-Brown et al. 2014; Kofler & Schlötterer 2014). Similar to Baldwin-Brown et al. (2014), we did not model allele frequency estimation errors caused for example by sequencing errors, limited read depth, read depth heterogeneity across the chromosome, or the number of sequenced individuals. We used PoPoolation2 (Kofler et al. 2011) to rescale allele counts for each biallelic position to a uniform read depth of 80.

### Experimental parameters

The purpose of this study is to test the influence of different experimental parameters on the power and resolution of secondary E&R. For this, we systematically varied the crossing scheme, population size, experimental duration, and number of replicates to assess the effect of these experimental parameters on the power and resolution of secondary E&R (Table 1). We use the term “experimental design” to describe a distinct set of simulated experimental parameters (e.g., crossing scheme=1:1_1f; population size=1,200 individuals; experimental duration=60 generations; 5 replicates).

**Table 1.** Simulation overview. 1:1_1f, 1:few, 1:many, dil:st, dil:mt: For each selected SNP, selection coefficients were uniformly sampled between 0.07 and 0.1. Dominance coefficients were randomly chosen to be either 0.5 (co-dominant) or 1 (dominant). 1:1_2f, 1:1_1f1nf: For each simulation, the selection coefficient of the target of interest was uniformly sampled between 0.07 and 0.1. In contrast to 1:1_1f, we simulated an additional selection target, which was either located on the focal haplotype (1:1_2f) or one non-focal haplotype (1:1_1f1nf). The selection coefficient of the additional selection target was uniformly sampled between 0.07 and the selection coefficient of the target of interest. We simulated all possible combinations of dominance coefficients for the two selected SNPs (500 simulations per combination).

| Crossing scheme | Population size | Generations | Replicates | Number of simulations |
| --- | --- | --- | --- | --- |
| *1:1_1f* | 300; 1,200; 4,800; 19,200 | 60; 20 | 5/30 | 2,000/100 |
| *1:1_2f* | 300; 1,200; 4,800; 19,200 | 60 | 5 | 2,000 |
| *1:1_1f1nf* | 300; 1,200; 4,800; 19,200 | 60 | 5 | 2,000 |
| *1:few* | 300 | 60 | 5/30 | 2,000/100 |
| *1:many* | 300 | 60; 20 | 5/30 | 2,000/100 |
| *dil:st* | 300 | 60 | 5/30 | 2,000/100 |
| *dil:mt* | 300; 1,200; 4,800; 19,200 | 60; 20 | 5/30 | 2,000/100 |

#### Crossing scheme

We simulated five different crossing schemes: 1:1 (versions 1f, 2f and 1f1nf), 1:few, 1:many, dil:st, and dil:mt (Figure 1B-F). In the **1:1_1f** crossing scheme (Figure 1B), two inbred founder lines are crossed at equal proportions. One inbred focal line carries a single target of selection – known from a primary E&R experiment – and is crossed with inbred non-focal lines, not carrying known adaptive alleles. We consider this a best-case scenario. Replicates are created by crossing the same focal line to different non-focal lines without known targets of selection. We used different non-focal lines as crossing partners to account for the possibility that in real experiments these lines may contain unidentified selected loci. By using different lines, the influence of selection targets present in a single non-focal line will be outweighed by the focal locus, which is present in every replicate (Figure 1B). To further explore the influence of unidentified selected loci, we simulated two more versions of the 1:1 crossing scheme with a more realistic genetic architecture (Figure S1). In these versions, either the focal line itself, or one of the non-focal lines carries one additional target of selection. We call these scenarios 2f, for a total of 2 selected loci in the focal line (Figure S1B), and 1f1nf, for one selected locus in the focal line and one selected locus in one of the non-focal lines (Figure S1C). For each simulation, we sampled the selection coefficient of the additional selection target uniformly between 0.07 and the selection coefficient of the selected SNP we consider for our analysis. We simulated all possible combinations of dominance coefficients for the two selected SNPs (0.5 – 0.5; 0.5 – 1; 1 – 0.5; 1 – 1). We ran 500 simulations for each of the 4 combinations of dominance coefficients for 2f and 1f1nf, respectively.

In the **1:few** crossing scheme (Figure 1C) the focal line is crossed with a pool of five non-focal lines that do not carry known beneficial alleles. The starting frequency of the focal line is 50 %, whereas each non-focal line has a starting frequency of 10 %. Each replicate consists of the same focal/non-focal line mixture. In the **1:many** crossing scheme (Figure 1D), the focal line is crossed with a pool of 99 non-focal lines. Replicates consist of the same mixture of lines, and the starting frequency of the focal line is 50 %.

It has been recently suggested, that “diluting” evolved populations of a primary E&R experiment with many non-adapted ancestral genotypes of the very same primary E&R and exposing the diluted populations to the same selection regime is a promising approach to validate selection candidates (Burny et al. 2020). However, we lack a systematic power assessment of such experiments with computer simulations. We thus included two “dilution” crossing schemes (**dil:mt, dil:st**) into our analysis. To evaluate the power of dilution crossing schemes, it is important to first simulate a primary E&R study. We chose to simulate a population consisting of 100 different founder lines, a population size of 300 individuals, 60 generations of adaptation and one replicate, which can then be diluted with non-adapted ancestral genotypes. SNPs in the primary E&R were tested for allele frequency change with the χ^2^ test.

The population of the **dil:st** crossing scheme carries only one beneficial SNP (**dil**:**st** = **dil**ution: **s**ingle **t**arget, Figure 1E). The starting allele frequency of this beneficial, focal SNP is sampled from the empirical starting allele frequency distribution of putatively selected alleles from a previous E&R study in which *D. simulans* populations adapted to a new temperature regime (mean starting frequency = 0.1) (Barghi et al. 2019). After simulating 60 generations of adaptation (primary E&R), 50 % of the evolved population is replaced by flies of the non-adapted ancestral founder population. After this dilution step, the secondary E&R was simulated under the exact same selection regime as in the primary E&R. A region with a strong selection signal from the primary E&R is chosen for validation in the dilution crossing schemes (Burny et al. 2020). Hence, we investigated a 1 Mb window, which is the previously reported median selected haplotype block length on chromosome-arm 2L (Barghi et al. 2019), around the SNP with the highest χ^2^ test-statistic in the primary E&R, in the secondary E&R.

The **dil:mt** (Figure 1F) crossing scheme has multiple selection targets (**dil**:**mt** = **dil**ution: **m**ultiple **t**argets; 16 selection targets on chromosome-arm 2L (Barghi et al. 2019)). Again, we investigate a 1 Mb window around the SNP with the highest χ^2^ test statistic in the primary E&R, in the secondary E&R. In case of multiple selection targets in the 1 Mb window, we consider the target that is closest to the SNP with the highest CMH test statistic in the 1Mb window of the secondary E&R in our analysis.

#### Population size

For two crossing schemes that are relatively easy to implement in empirical studies 1:1 (1f, 2f and 1f1nf) and dil:mt we performed simulations (experimental duration = 60 generations; 5 replicates) with independently sampled selection targets for population sizes of 300; 1,200; 4,800; and 19,200 individuals per replicate to test for the effect of the population size on the power and resolution of secondary E&R (Table 1). All other crossing schemes were evaluated at a population size of 300 individuals.

#### Experimental duration

All crossing schemes were evaluated after 60 generations. To explore the possibility that shorter experiments with less than 60 generations may already be sufficient to achieve satisfactory power, we analyzed the simulations for 1:1 (without 2f and 1f1nf) and dil:mt crossing schemes also already after 20 generations (Table 1).

#### Number of replicates

To assess the impact of the number of replicates on power of secondary E&R, we conducted for each crossing scheme (1:1 without 2f and 1f1nf) 100 additional simulations (population size= 300 individuals; experimental duration = 60 generations) with 30 replicates (Table 1).

### Statistical analysis

All statistical analyses were performed using the R statistical computing environment (v3.5.3) (R Core Team 3.5.3 2019).

#### Power

For each simulation, we tested all SNPs on chromosome-arm 2L for allele frequency increase between the start and the end of the simulated secondary E&R using the Cochran-Mantel-Haenszel-test (CMH-test, implemented in the R package poolSeq (v0.3.2) (Taus et al. 2017)). The CMH-test allows to test for independence of matched data – e.g. allele counts of replicated ancestral and evolved populations (Agresti & Kateri 2011). We ranked SNPs based on their CMH test statistic using the dense ranking method in the R package data.table (v.1.12-8) (Dowle & Srinivasan 2019). In dense ranking, SNPs with identical test statistics receive the same rank, and the following SNP is assigned the immediately following rank.

Based on this ranking, we used two different approaches to classify simulations being either successful, or unsuccessful. First, we only considered a simulation to be successful if the true target of selection (i.e., the focal SNP with a selection coefficient > 0) was the SNP with the highest test statistic (success-A). In a second analysis step, we classified a simulation as success, if the focal target of selection was not more than 100 SNPs away from the SNP with the highest test statistic (success-B, Figure 2). We acknowledge that success-B depends on the maximum distance allowed between the true target of selection and the SNP with the highest CMH test statistic (Figure 2). However, varying the maximum distance threshold did not alter the relative performance of different experimental designs (data not shown). The power of an experimental design is defined as the proportion of simulations that were able to detect the true target of selection.

**Figure 2.**
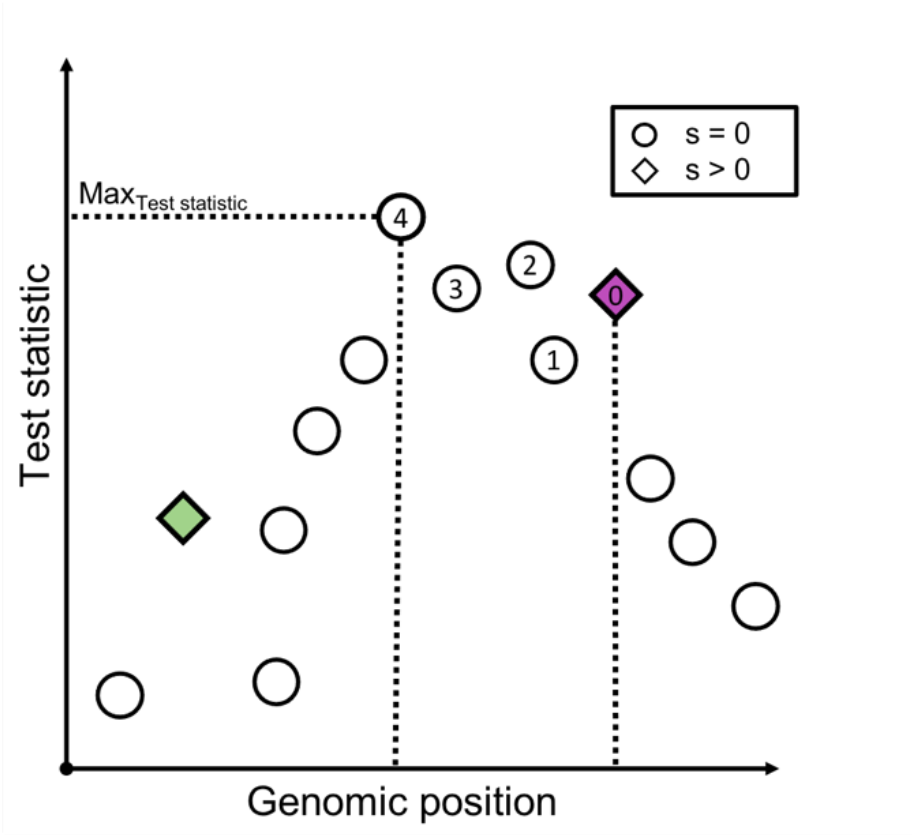
Schematic overview of the definition of success-B in a secondary Evolve and Resequence simulation. All SNPs (neutral = circle, beneficial = diamond) are tested for an allele frequency change with the Cochran-Mantel-Haenszel-test, and are ranked based on their test statistic (y-axis). If the focal selection target (purple diamond; one additional selected SNP is shown as green diamond - for crossing scheme 1:1_2f, 1:1_1fnf, and dil:mt) is less than 100 SNPs away from the SNP with the highest test statistic, the simulation run is deemed a success. In the example depicted, the distance in number of SNPs between the focal target of selection, and the SNP with the highest test statistic is 4, and the simulation is classified as success.

#### Resolution

For simulations where the selection target was detected (success-B), we determined the resolution of fine mapping of the selection target by counting the number of SNPs between the true selection target, and the SNP with the highest CMH test statistic.

#### Assessment of experimental and population genetic parameters

We used logistic regressions with a binomial error structure (*ε*) and a logit link function (Baayen 2008) to test how different experimental and population genetic parameters affect success and failure to identify targets of selection, i.e. success (Y) is treated as a binary response encoded in 0 (failure) and 1 (success) of secondary E&R.

We fitted three different models with R function *glm* with *µ* being the overall mean per model: Model 1 includes only the two crossing schemes 1:1_1f and dil:mt for which we also varied population size. Model 2 includes the three versions of crossing scheme 1:1 (1f, 2f and 1f1nf) at varying population sizes: 1f with only one positively selected SNP in the focal line; version 2f with two selected SNPs in the focal line; and version 1f1nf with one selected SNP in the focal line and one selected SNP in one non-focal line. Model 3 includes all 5 simulated crossing schemes (1:1 without 2f and 1f1nf) at a constant population size of 300 individuals.

Prior to model fitting, the two covariates selection coefficient and mean starting allele frequency (not applicable to model 2 as starting allele frequency is always 50%) were multiplied by 100, and z-transformed to a mean of zero and standard deviation of one for easier interpretable estimates (Schielzeth 2010).

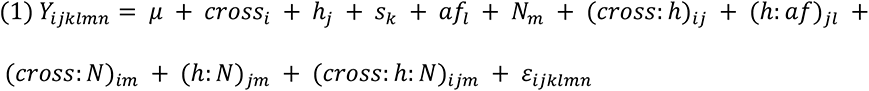

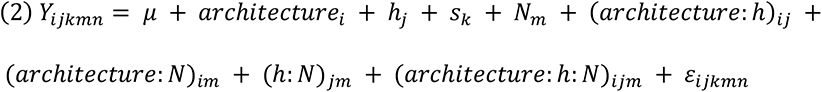

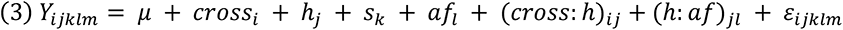

Model 1 (equation (1)) contained (i) five explanatory variables as main effects; crossing scheme (*cross_i_*), a fixed categorical effect with 2 levels - 1:1_1f and dil:mt; a fixed categorical effect of dominance coefficient (*h_j_*) with levels 0.5 and 1; selection coefficient (*s_k_*) and mean starting allele frequency (*af_l_*) over replicated populations (both as continuous covariate); and population size (*N_m_*), a fixed categorical effect with 4 levels - 300; 1,200; 4,800; 19,200, (ii) an interaction term between dominance coefficient and mean starting allele frequency ((*h*:*af*)*_jl_*), (iii) and a triple interaction between crossing scheme, dominance coefficient, and population size ((*cross:h:N*)*_ijm_*), and all pairwise interaction terms of effects involved in the triple interaction term ((*cross:h*)*_ij_*; (*cross:N*)*_im_*; (*h:N*)*_jm_*). We included interaction terms into the model that have population genetic interpretations.

Data analyzed with model 1 contained 16,000 observations, namely 2,000 independent simulation runs for each crossing scheme (1:1_1f, dil:mt), and each of the four different population sizes (300; 1,200; 4,800; 19,200) (Table 1). To avoid potential bias introduced by specific haplotypes being sampled, we randomly chose 4 sets of focal/non-focal founder lines for the 1:1_1f crossing scheme and performed 500 simulation runs per set. The chosen set of founder lines did not have a significant effect on the success of secondary E&R and is thus not included in the final model (likelihood ratio test (LRT) full-reduced model comparison; data not shown).

Model 2 (equation (2)) contained (i) four explanatory variables as main effects; a fixed categorical effect “architecture” (*architecture_i_*) that describes the version of the 1:1 crossing scheme in combination with the dominance coefficient of the additional selected target (if present) resulting in 5 levels: 1f; 2f_h05; 2f_h1; 1f1nf_h05; 1f1nf_h. Model 2 further contained a fixed categorical effect of dominance coefficient for the focal SNP (*h_j_*) with levels 0.5 and 1; selection coefficient (*s_k_*) as continuous covariate; and population size (*N_m_*), a fixed categorical effect with 4 levels: 300; 1,200; 4,800; 19,200, (ii) a triple interaction between architecture, dominance coefficient, and population size ((*architecture*:*h*:*N*)*_ijm_*), and all pairwise interaction terms of effects involved in the triple interaction term ((*architecture*:*N*)*_im_*; (*h*:*N*)*_jm_*).

Data analyzed with model 2 contained 24,000 observations, namely 2,000 independent simulation runs for each version of the 1:1 crossing scheme (1f, 2f, 1f1nf), and four different population sizes (300; 1,200; 4,800; 19,200 individuals). Because model 1 showed that the chosen set of founder lines did not have a significant effect on the power of secondary E&R, we simulated only one set of focal/non-focal founder lines.

Model 3 (equation (3)) contained (i) four explanatory variables; crossing scheme (*croos_i_*), a fixed categorical effect with 5 levels: 1:1_1f; 1:few; 1:many; dil:st; dil:mt, dominance coefficient (*h_j_*), selection coefficient (*s_k_*), and mean starting allele frequency (*af_l_*) over replicated populations as main effects, as described for model 1, (ii) an interaction term between crossing scheme and dominance coefficient ((*cross:h*)*_ij_*), (iii) and an interaction term between dominance coefficient and mean starting allele frequency ((*h*:*af*)*_jl_*).

We analyzed 10,000 samples (2,000 independent simulation runs for each crossing scheme) (Table **1**). For crossing schemes using only few different founder lines (1:1_1f, 1:few), the simulation runs are based on 4 randomly chosen sets of founder lines each (500 simulation runs per set). As in model 1, the set of founder lines does not have a significant effect on secondary E&R success and is thus not included in the final model (LRT full-reduced model comparison; data not shown). We performed additional analysis with 500 observations (100 independent simulation runs for each crossing scheme) to determine the influence of the number of replicates on the power of secondary E&R (Table 1).

We performed all diagnostic checks required for logistic regression. Absence of collinearity was confirmed by computing the generalized Variance Inflation Factors (Fox & Monette 1992) using function *vif* in R package *car* (v3.0-8 (Fox & Weisberg 2019)). Model stability was checked with the R function *dfbeta*. For visualization, linear predictors (LP) were back-transformed to success probabilities using the inverse logit transformation: 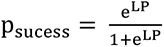. 95% confidence intervals of the fitted values were investigated with the R function *predict.glm*. Significance of single explanatory variables was tested with a Type II ANOVA using function *Anova* in R package car (Fox & Weisberg 2019) and are provided in the Supplement. Significance of explanatory variables including all their modeled interactions was tested with a likelihood ratio test comparing the full model with a nested reduced model with the same structure as the full model, but lacking the assessed explanatory variable (and its interactions). Significance is declared at an alpha cut-off of 5%. We used Nagelkerke’s R^2^ index (Nagelkerke 1991) to calculate the improvement of each model parameter upon the prediction of a reduced model.

We observed two cases where a combination of explanatory variables resulted in complete separation of data points (Figure 5A (dominance coefficient: 0.5, population size: 1,200, architecture: 2f_h05), Figure S8A (dominance coefficient: 0.5, crossing scheme: dil:mt)). To obtain interpretable model estimates, we added one pseudo-observation with the missing response (success) to the data for each of these two cases.

## Data availability

Information regarding the accessibility of raw sequence reads, phased haplotypes of the ancestral *D. simulans* haplotypes as well as MimicrEE2 ready text files for the *D. simulans* recombination map (file used in this project: Dsim_recombination_map_LOESS_100kb_1.txt) can be found in (Howie et al. 2019). MimicrEE2 ready input files of the different experimental designs, the simulated selection regimes, processed simulation results and all scripts that are necessary to reproduce the results are available at SourceForge (https://sourceforge.net/projects/secondary-e-r-sim/files/).

## Results

We used forward simulations to assess the influence of different experimental and population genetic parameters, more specifically crossing scheme, population size, dominance coefficient, selection strength, and mean starting allele frequency on the success to detect and fine map selection targets in secondary E&R experiments (Figure 1A). We used a Cochran-Mantel-Haenszel (CMH) test to identify SNPs rising in allele frequency (number of SNPs see Table S1). The CMH test allows to test for independence of matched categorical data (Agresti & Kateri 2011), and compares favorably to other statistical methods in reliably identifying possible targets of selection in E&R setups (Vlachos et al. 2019). A simulation run was considered successful, if the true target of selection was the SNP with the highest CMH test statistic (success-A). In a second analysis step, we considered simulation runs as successful, if the true target of selection was not more than 100 SNPs away from the SNP with the highest CMH test statistic (success-B, Figure 2). We used logistic regression to assess which experimental and population genetic parameters have a significant effect on the success of secondary E&R.

First, we evaluated two crossing schemes that can be easily implemented in empirical studies, 1:1_1f (Figure 1B) and dil:mt (Figure 1F). The 1:1_1f crossing scheme (Figure 1B) is based on founder lines only (a founder line is an inbred isofemale line homozygous for one ancestral haplotype). Using information about the selected haplotype from a primary E&R study, it is possible to determine which founder lines carry a selection target (Barghi et al. 2019). Note, this requires focal founder lines to be sequenced. Crossing a focal founder line with the selected haplotype with a non-focal line without known selection targets offers the advantage of reducing the number of selection targets dramatically. Using different non-focal lines without known strong selection targets in each replicate reduces the potential of consistent confounding effects of unidentified selection targets outside the region of interest – a signal the CMH test is particularly sensitive to as it scans for consistent allele frequency changes across replicates.

Dil:mt (Figure 1F) represents an entirely different approach. Dil:mt has multiple targets of selection at different frequencies and is probably the simulated crossing scheme with the most straight forward empirical implementation (Barghi et al. 2019; Burny et al. 2020). It is based on a “dilution” approach, where evolved individuals from a primary E&R experiment are crossed to the non-adapted ancestral founder population from the same primary E&R (Burny et al. 2020). In contrast to 1:1_1f, a dil:mt crossing scheme requires both – ancestral and evolved – populations of the primary E&R, but relatively limited information about selection targets on individual founder lines.

To evaluate the 1:1_1f and dil:mt crossing scheme, we simulated secondary E&R consisting of 5 replicated populations with constant population size (300; 1,200; 4,800, or 19,200 individuals per replicate) that evolve for 60 generations. We simulated positively selected SNPs (selection coefficient is uniformly sampled between 0.07 and 0.1) that were randomly chosen with equal probability to be either co-dominant (h=0.5) or dominant (h=1). We used logistic regression to assess the effects of model parameters on secondary E&R success (Model 1, *Assessment of experimental and population genetic parameters* in Material & Methods). While selection strength (LRT full-reduced null model comparison (Model 1): χ^2^ = 67.4, df = 1, p<0.001 (success-A); χ^2^ = 73.4, df = 1, p<0.001 (success-B)), dominance coefficient (LRT full-reduced model comparison (Model 1): χ^2^ = 733.4, df = 9, p<0.001 (success-A); χ^2^ = 714.5, df = 9, p<0.001 (success-B)), and mean starting allele frequency (LRT full-reduced model comparison (Model 1): χ^2^ = 22.4, df = 2, p<0.001 (success-A); χ^2^ = 17.4, df = 2, p<0.001 (success-B)) all have a significant effect on the power of secondary E&R (Table S2), our analysis reveals that crossing scheme and population size have by far the strongest influence on secondary E&R success. Both the crossing scheme (LRT full-reduced model comparison (Model 1): χ^2^ = 3510.1, df = 8, p<0.001 (success-A); χ^2^ = 3211.3, df = 8, p<0.001 (success-B)); and the population size (LRT full-reduced model comparison (Model 1): χ^2^ = 2142.9, df = 12, p<0.001 (success-A); χ^2^ = 2633.4, df = 12, p<0.001 (success-B)) have a significant effect on the success of secondary E&R, and are the only parameters with a Nagelkerke’s R^2^ index above 0.2 (Table S3).

While a 1:1_1f crossing scheme results in higher power values than the dil:mt crossing scheme independently of the population size, the difference to the dil:mt crossing scheme is more pronounced in larger experimental populations (Figure 3). This does not hold only for the power, but also for the resolution (= the distance in SNPs between the true target of selection, and the SNP with the highest CMH test statistic, Figure 4). A potential disadvantage of the 1:1_1f crossing scheme is that the low number of different founder lines (n=6, Figure 1B) causes more linkage disequilibrium, indicated by the number of neighboring SNPs with identical test statistics (=ties) and thus broadens signatures compared to dil:mt crossing scheme that has more founder lines (n=100) (Figure S2). However, this is outweighed by superior power of the 1:1_1f crossing scheme at every population size investigated (Figure 3-4).

**Figure 3.**
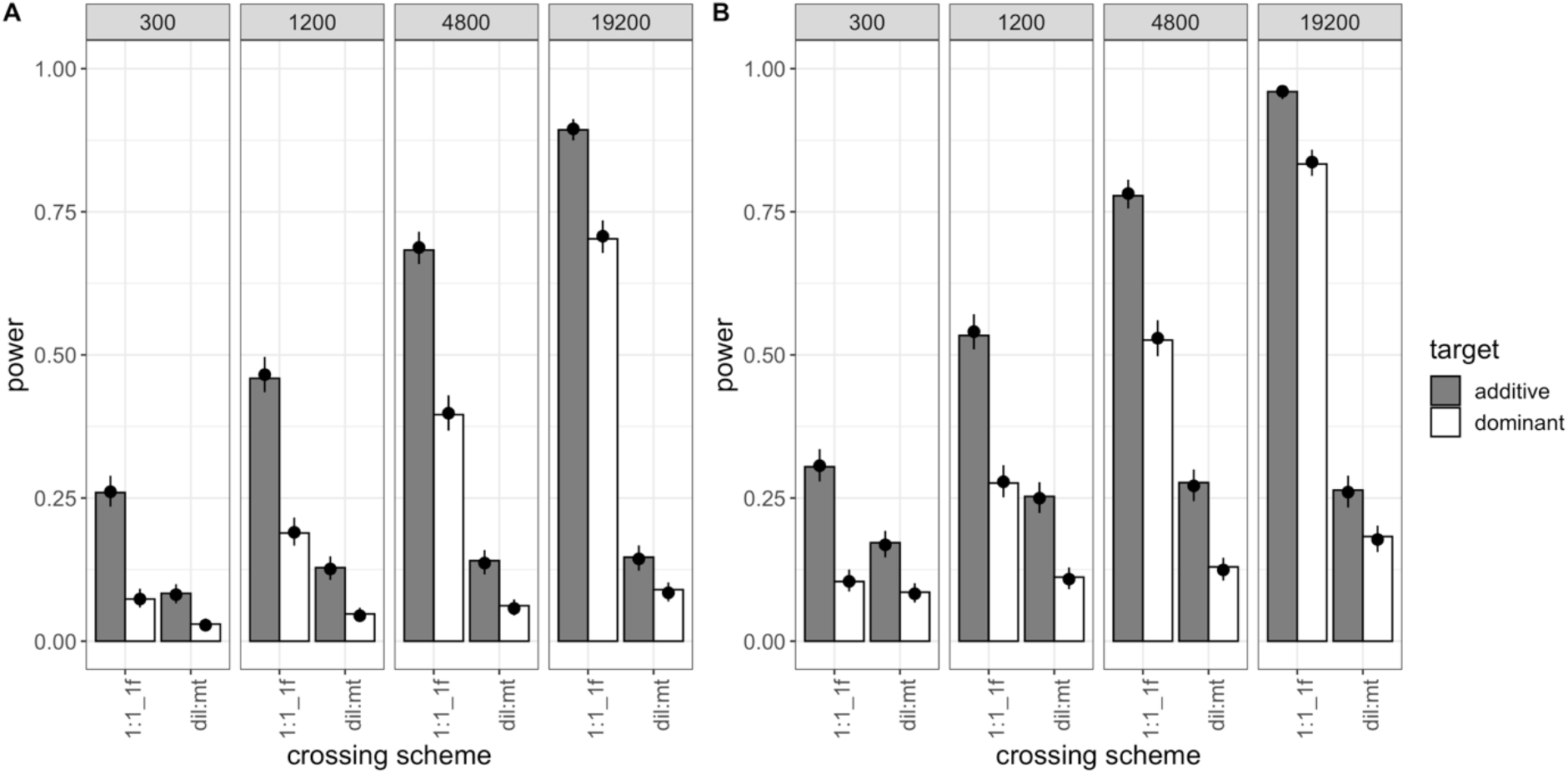
Power of the 1:1_1f and dil:mt crossing scheme at different population sizes (2,000 simulations/experimental design). Bars show the power (i.e., the proportion of successful simulations) separately for each combination of crossing scheme (1:1_1f, dil:mt), population size (300; 1,200; 4,800; 19,200 individuals), and dominance coefficient (additive in grey, dominant in white). The dots with error bars display the estimate from the fitted model (Model 1) and its 95 % confidence interval. For the model fit, the selection coefficient was fixed to its global average, and combination-specific average starting allele frequencies were used. (A) shows the results for success-A (= selection target is the SNP with the highest Cochran-Mantel-Haenszel (CMH) test statistic), (B) shows the results for success-B (= selection target is not more than 100 SNPs away from the SNP with the highest CMH test statistic).

**Figure 4.**
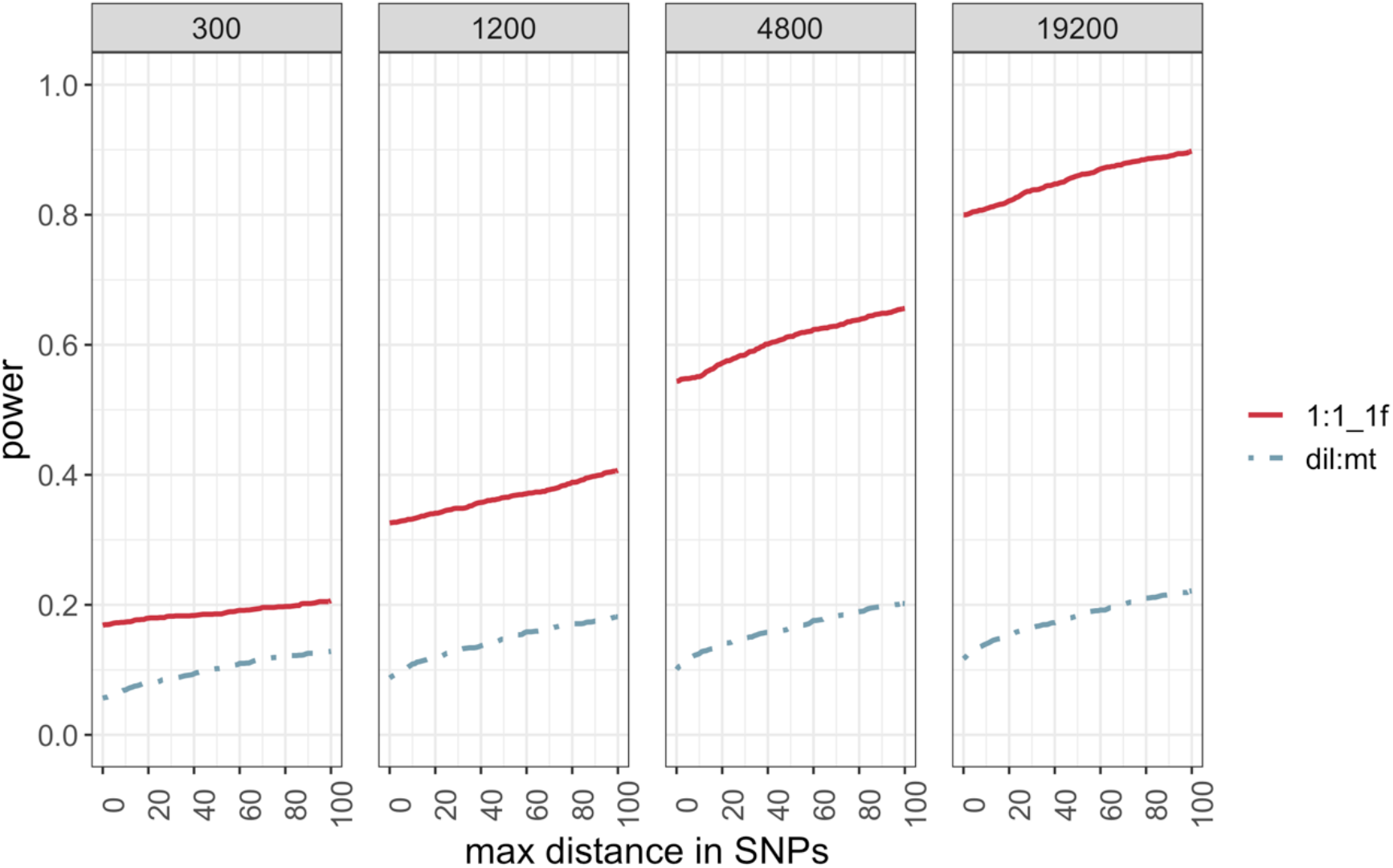
Resolution of the 1:1_f and dil:mt crossing scheme at different population sizes. Proportion of simulations (y-axis) that do not exceed a maximum distance in SNPs (x-axis) between the SNP with the highest Cochran-Mantel-Haenszel test statistic and the true target of selection. Each panel shows the result for one simulated population size.

### Experimental duration

Our analysis showed that after 60 generations of adaptation, only experimental designs with a population size of at least 1,200 individuals have more power than 50 % (Figure 3). The 1:1_1f crossing scheme clearly outperforms dil:mt regardless of the population size, and reaches power above 75% only with a population size of at least 4,800 individuals. However, the maintenance of 5 replicates for 60 generations is labor intensive and can be quite time consuming for most sexual organisms. We explored the possibility that shorter experiments with less than 60 generations may already be sufficient to achieve satisfactory power values. We thus reanalyzed the power of the 1:1_1f, and dil:mt crossing scheme after 20 generations of adaptation (Table 1).

Consistent with previous computer simulation studies (Baldwin-Brown et al. 2014; Kofler & Schlötterer 2014; Kessner & Novembre 2015) and empirical results (Langmüller & Schlötterer 2020), we observe reduced power for experimental designs with shorter experimental duration (Figure S3). Consistent with the results after 60 generations, the crossing scheme (LRT full-reduced model comparison (Model 1): χ^2^ = 2672.7, df = 8, p<0.001 (success-A); χ^2^ = 2234.4, df = 8, p<0.001 (success-B)) and the population size (LRT full-reduced model comparison (Model 1): χ^2^ = 3176.4, df = 12, p<0.001 (success-A); χ^2^ = 3375.3, df = 12, p<0.001 (success-B)) have the biggest effects on the success of secondary E&R (TableS2-3).

In contrast to our analysis after 60 generations of adaptation, the dil:mt crossing scheme results in higher power than the 1:1_1f crossing scheme for populations with the smallest simulated population size (300 individuals) after 20 generations of adaptation (Figure S4). A possible explanation for this is that the dil:mt crossing scheme has multiple beneficial targets. Because we simulate additive selection, linked selection targets can act synergistically and increase the frequency of the focal SNP over shorter time scales. This phenomenon of pronounced allele frequency increase due to linked selection will be especially important if the population size is small (i.e., drift is not neglectable), and the experimental duration is short. This is reflected in a higher Nagelkerke’s R^2^ index for population size in secondary E&R with an experimental duration of 20 generations compared to 60 generations of adaptation (Table S3).

With increasing population size and thus, reduced genetic drift, the 1:1_1f crossing scheme outcompetes dil:mt, as already seen in our analysis after 60 generations (Figure S3-4). Although shorter experimental duration reduces power, our analysis after 20 generations of adaptation highlights that if the maintenance of experimental populations for many generations is not feasible, shorter experiments can achieve similar power if they are maintained at larger population sizes (Figure 3, Figure S3).

### Additional target of selection in the 1:1 crossing scheme

In contrast to dil:mt, the 1:1_1f crossing scheme harbors only one target of selection. We simulated two additional versions of the 1:1 crossing scheme to investigate how one additional target of selection influences the power of secondary E&R (Figure S1; Model 2, *Assessment of experimental and population genetic parameters* in Material & Methods). We observed that one additional beneficial SNP has a significant effect on the power of secondary E&R using the 1:1 crossing scheme (LRT full-reduced model comparison (Model 2): χ^2^ = 1727.7, df = 32, p<0.001 (success-A); χ^2^ = 2588.42, df =32, p<0.001 (success-B); Table S4-5). This significant effect is mainly driven by one scenario: when the focal line harbors two targets of selection with the target of interest being dominant and the additional target being co-dominant (Figure 5) the power to detect the target of interest is close to zero. The reason is that with a starting frequency of 50 %, co-dominant alleles respond more to strong selection than dominant alleles, because non-favored alleles are masked by high frequency dominant alleles (Figure S5). This differential behavior is particularly pronounced for high allele frequencies. Note that our definitions of success do not use prior information on the location of the focal SNP of interest. We propose that a substantial fraction of the power can be recovered if the analysis is restricted to the approximate location of the focal target determined in the primary E&R study. On the other hand, for additive targets of interest one additional target of selection hardly reduces the power of the 1:1 crossing scheme, regardless of the dominance coefficient of the additional target and whether the additional target is positioned on the focal or one non-focal haplotype (Figure 5). Overall, our results show that the 1:1 crossing scheme still outperforms dil:mt even in the presence of one additional target of selection (Figure 3, Figure 5). For the remaining analysis we will focus on a 1:1_1f crossing scheme with only one target of selection.

**Figure 5.**
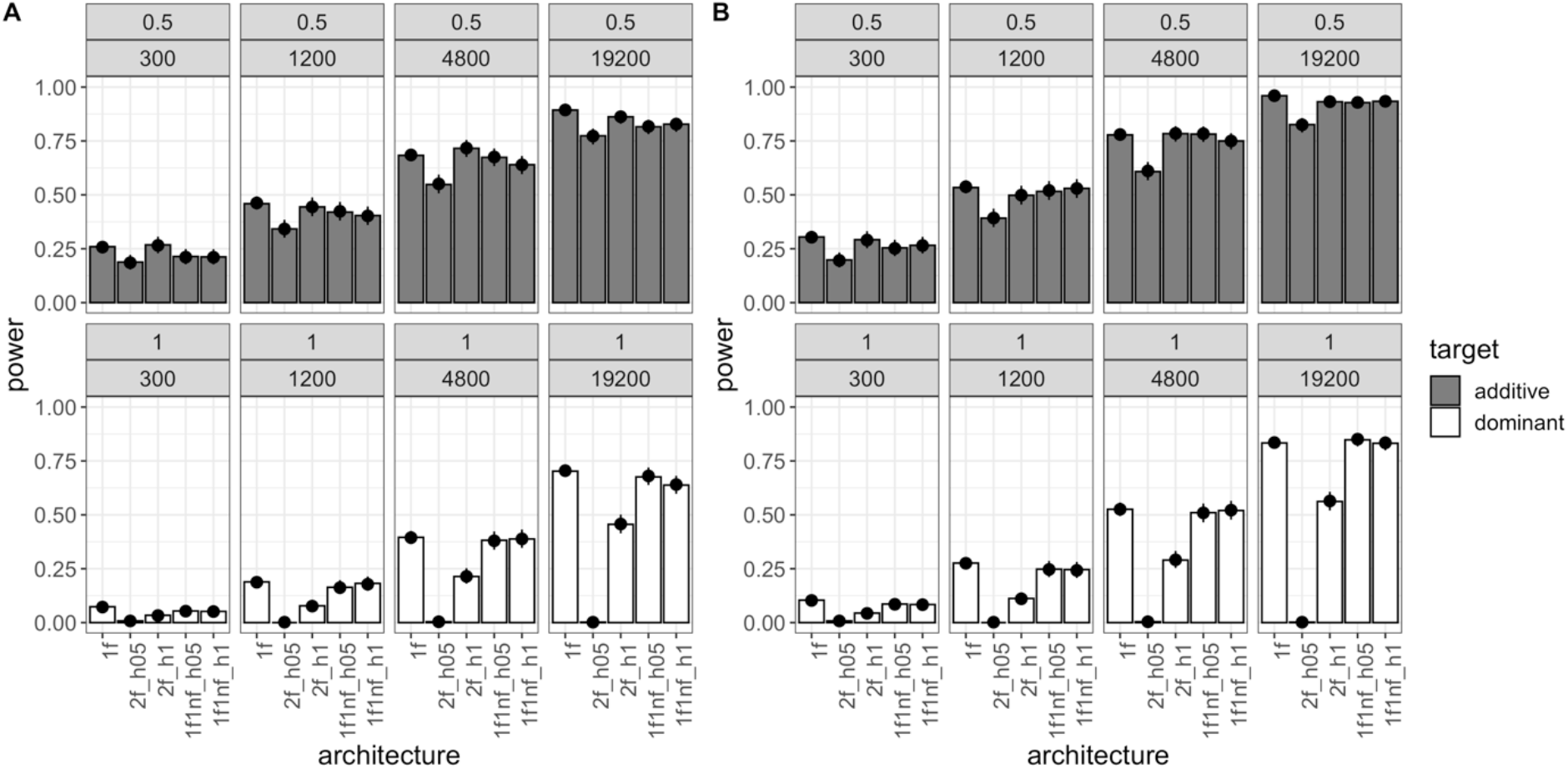
Power of the different 1:1 crossing schemes at different population sizes (Model 2) (2,000 simulations/experimental design) after 60 generations of adaptation. Bars show the power (i.e., proportion of successful simulations) separately for each combination of 1:1 crossing scheme version with the dominance coefficient of the additional target of selection: 1f, only one target of selection on the focal haplotype; 2f_h05/2f_h1, 2 targets of selection on the focal haplotype; 1f1nf_h05/1f1nf_h1, 1 target of selection on the focal haplotype and one on one non-focal haplotype. Columns depict different population sizes (300; 1,200; 4,800; 19,200 individuals), and rows show different dominance coefficients of the target of interest (additive in top row, dominant in bottom row). The dots with error bars display the model fit (Model 2) and its 95 % confidence interval. For the model fit, the selection coefficient was fixed to its global average. (A) shows the results for success-A (= selection target is the SNP with the highest Cochran-Mantel-Haenszel (CMH) test statistic) (B) shows the results for success-B (= selection target is not more than 100 SNPs away from the SNP with the highest CMH test statistic).

### Alternative crossing schemes

Since large population sizes can be challenging to maintain, we also evaluated additional crossing schemes which may perform better even for smaller population sizes (Model 3, *Assessment of experimental and population genetic parameters* in Material & Methods). 1:few (Figure 1C) is a modification of the 1:1_1f crossing scheme, which combines all non-focal lines in each replicate - this provides a consistent genetic composition in each replicate. The 1:many crossing scheme (Figure 1D) uses 1 focal, and 99 non-focal lines without selection target. We reasoned that the larger number of segregating variants (970,466 SNPs compared to max 527,771 SNPs in 1:1 crossing scheme, Table S1) potentially provides a higher mapping resolution. Finally, we modified the dilution crossing scheme by simulating only one selection target in the founder population (dil:st, Figure 1E). Hence, no additional selection targets can create potentially confounding selection signatures. For each experimental design, we simulated a population size of 300 individuals, 60 generations of adaptation, and 5 replicates (Table 1).

Regardless of the experimental design, the simulated focal selection targets experienced a pronounced allele frequency increase (Figure S6). In “dilution” crossing schemes (dil:st, dil:mt; Figure 1E-F) the frequency trajectories of the focal SNPs were highly variable because of the heterogeneous starting frequency. The crossing schemes where a single focal line with a starting frequency of 50 % carries the beneficial allele typically had a superior performance than dilution crossing schemes (Figure 6). Notably, the significant influence of crossing scheme on secondary E&R success (LRT full-reduced model comparison (Model 3): χ^2^ = 295.9, df = 8, p<0.001 (success-A); χ^2^ = 190.23, df =8, p<0.001 (success-B)) (Table S6-7) cannot be explained by a loss of the focal SNP in dilution crossing schemes, which occurred in less than 1 % of the simulations.

**Figure 6.**
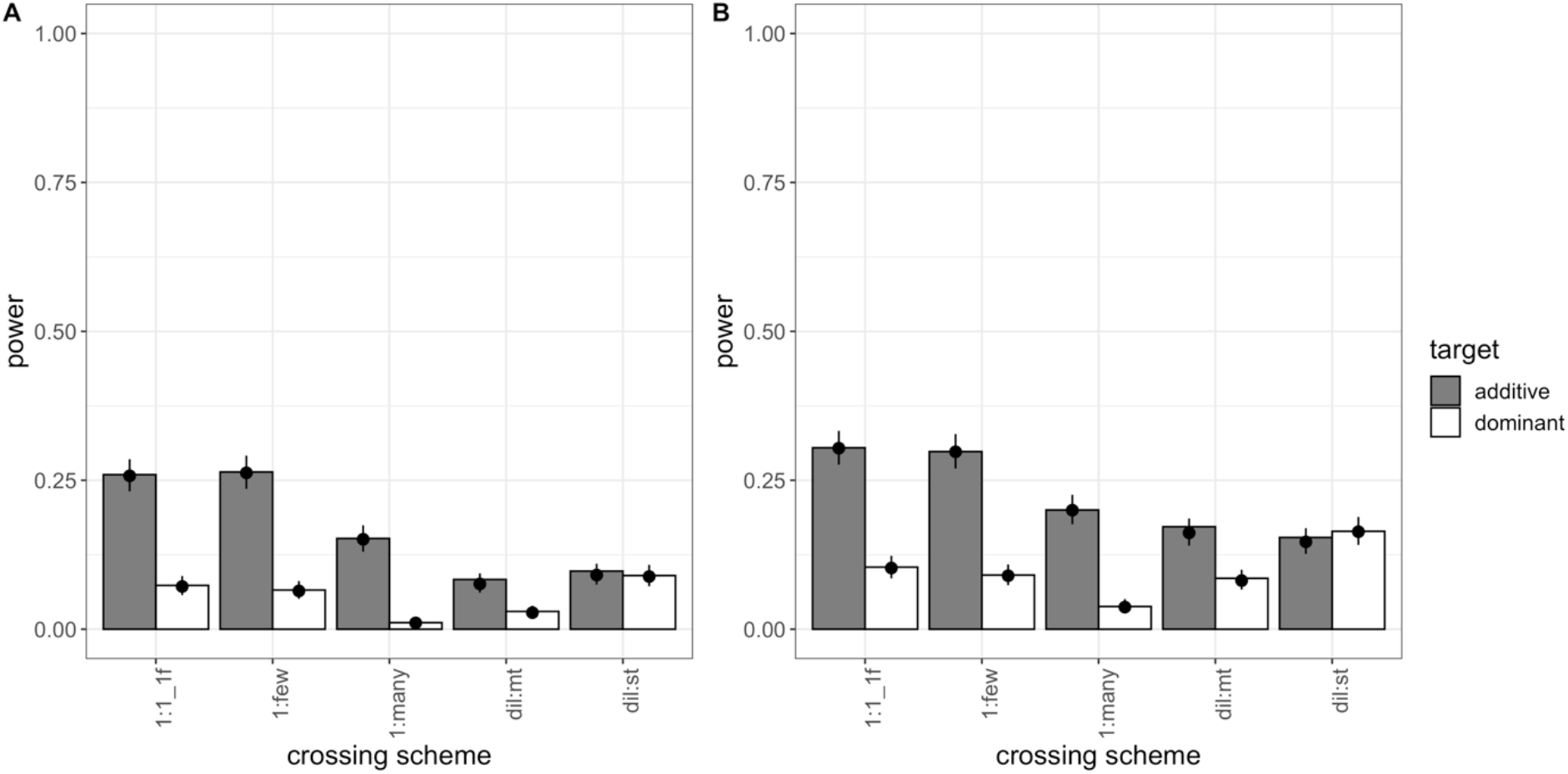
Power of five different crossing schemes (population size = 300 individuals; 2,000 simulations/experimental design). Bars show the power (i.e., proportion of successful simulations) separately for each combination of crossing scheme and dominance coefficient (additive in grey; dominant in white). The dots with error bars display the estimate from the fitted model (Model 3) and its 95 % confidence interval. For the model fit, the selection coefficient was fixed to its global average, and combination-specific average starting allele frequencies were used. (A) shows the results for success-A (= selection target is the SNP with the highest Cochran-Mantel-Haenszel (CMH) test statistic), (B) shows the results for success-B (= selection target is not more than 100 SNPs away from the SNP with the highest CMH test statistic).

We found that the dil:st crossing scheme is still inferior to the 1:1_1f crossing scheme for additive loci (Figure 6). Also, the modifications of the 1:1_1f crossing scheme (1:few, 1:many) do not provide a substantial improvement (Figure 6). Surprisingly, the 1:many crossing scheme performed very poorly. This may be at least partly attributed to the larger number of SNPs/kb, which will affect the SNP-based success rate (success-B, Figure 2). Given that the three additional crossing schemes do not provide a clear advantage (Figure 6-7, Figure S7), and are not easier to execute experimentally, we did not evaluate them with larger population sizes.

**Figure 7.**
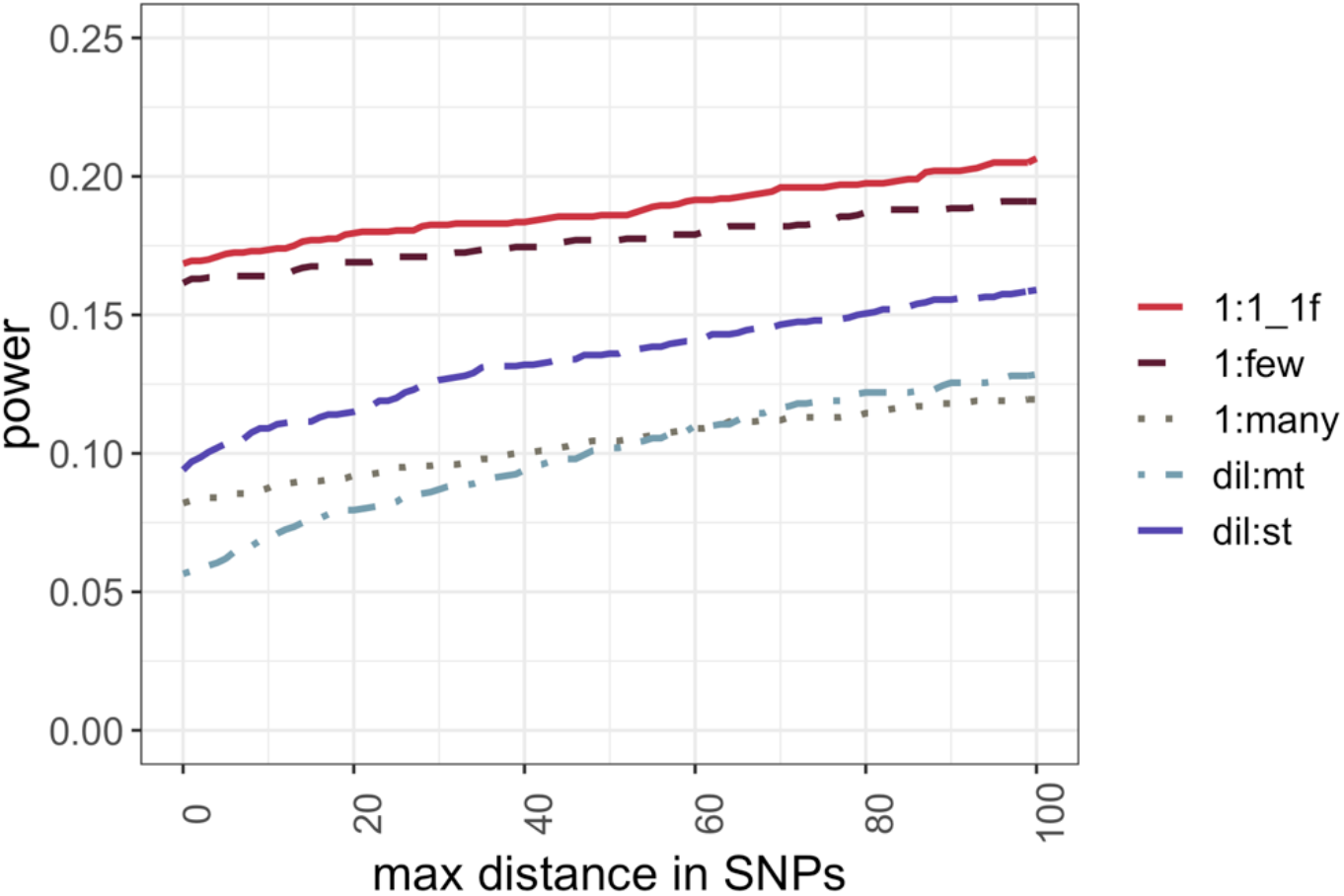
Resolution of five different crossing schemes. Proportion of simulations (y-axis) that do not exceed a maximum distance in SNPs (x-axis) between the SNP with the highest Cochran-Mantel-Haenszel test statistic and the true target of selection.

### Number of replicates

Given the superior performance of experimental designs with larger population sizes (Figure 3), we were also interested whether more replicates with a smaller population size may provide further improvements. We performed additional simulations for all five different crossing schemes with a population size of 300 individuals per replicate, and 30 replicates (Table 1). Consistent with other simulation studies (Kofler & Schlötterer 2014; Kessner & Novembre 2015), more replicates result in increased power (Figure S8). The power of secondary E&R with 30 replicates (Table S6) is only significantly affected by the crossing scheme considering all modeled interactions (LRT full-reduced model comparison (Model 3): χ^2^ = 31.467, df = 8, p<0.001 (success-A); χ^2^ = 16.692, df = 8, p=0.033 (success-B); Table S7). The 1:1_1f crossing scheme with 30 replicates also has the highest resolution (Figure S9A). As expected, the impact of linkage disequilibrium, indicated by the number of ties, is dramatically reduced for all crossing schemes when more replicates are simulated (Figure S9B compared to Figure S7). However, similar improvements as seen with 30 replicates can be achieved if 5 replicates of the 1:1_1f crossing scheme are used with a population size of 1,200 individuals (Figure 3). Thus, the same power can be achieved while maintaining about 33 % fewer individuals.

### Dominance coefficient, Selection coefficient, and mean starting allele frequency

As expected, dominance coefficient, selection coefficient, and average starting allele frequency have a significant effect on the success of secondary E&R in all simulation scenarios, except for data with 30 replicates. We observed that selection targets with a higher selection coefficient and/or a higher mean starting allele frequency are easier to fine map. The lack of significant effects of these parameters on secondary E&R success for the dataset with 30 replicates can probably be explained by the smaller number of conducted simulations (100 for 30 replicates and 2,000 for 5 replicates, Table 1). After crossing scheme and population size, the dominance coefficient is the parameter with the highest Nagelkerke’s R^2^ index in our analysis, with additive loci being easier to fine map. This is caused by the fact that heterozygotes and target homozygotes with a dominant beneficial SNP have the same fitness, resulting in less efficient selection that becomes especially apparent if the selected allele has already reached a high frequency in the population, and non-selected allele homozygotes become rare (indicated by a significant negative effect of the interaction term between dominance coefficient and mean starting allele frequency in most of our statistical models).

## Discussion

This work was inspired by the difficulty of most E&R studies with sexually reproducing organisms to pinpoint selection targets, mostly due to numerous neutral hitchhikers resulting in large haplotype blocks. Secondary E&R – a follow-up EE validating putative selection targets of a primary E&R under an identical selection regime – has been recently suggested as experimental approach for selection target confirmation (Burny et al. 2020). We used extensive computer simulations to evaluate how experimental and population genetics parameter shape the power and resolution of secondary E&R. As expected, dominance coefficient, selection strength, and mean starting allele frequency of the selected target all have a significant effect on the success of E&R, where dominant selection targets at high frequency are particularly challenging to detect. However, population size and crossing scheme emerged as the most influential parameters in our analysis.

### The crossing scheme has a pronounced effect on secondary E&R success

We show that a simple crossing scheme, which only requires that (a subset of) the founder lines are sequenced and that founder lines with and without the selection target of interest can be distinguished, has the best power and resolution of the five crossing schemes tested. The 1:1 crossing scheme is particularly well-suited when many selection targets are detected in the evolved populations of the primary E&R, because it uses only a subset of the lines from the non-adapted ancestral founder population of the primary E&R study. This reduces potential confounding effects between the selection target of interest and other adaptive loci in two ways. First, non-adapted ancestral founder haplotypes will harbor on average less selection targets than evolved haplotypes that can acquire multiple selection targets through recombination events during the primary E&R (Otte & Schlötterer 2021). Because evolved haplotypes will often harbor multiple selection targets, beneficial alleles will not propagate independently from each other, which makes it challenging to fine-map single selection targets. Second, the total number of potential beneficial alleles in a 1:1 crossing scheme is deliberately reduced by picking only a subset of the founder lines of the primary E&R, which facilitates fine mapping of one particular selection target of interest with secondary E&R.

We would like to point out that in our study “crossing schemes” are defined such that they do not only differ in the actual crossing procedure, but also in the number and starting allele frequency distribution of simulated selection targets (e.g., 1:1_1f vs. dil:mt). Additional beneficial alleles can without doubt have an impact on the power of secondary E&R. While the 1:1 crossing scheme still outperforms dil:mt in the presence of one additional selection target, the relative performance of crossing schemes might change given an even more complex underlying genetic architecture. Furthermore, we focused in this study on the assessment of experimental and population genetic parameters given a directional selection regime. Future research on the potential of secondary E&R to fine map selection targets under a complex/polygenic genetic architecture is needed, such as for example a quantitative trait under stabilizing selection that experienced a recent shift in trait optimum.

### Sufficient E&R power requires large population sizes

Our analyses also indicate that rather large population sizes are crucial to identify the causative variant with sufficient confidence using a secondary E&R approach. This observation explains why an empirical secondary E&R study in *D. simulans* with a population size of 1,250 flies per replicated population failed to narrow down a pronounced candidate region from a primary E&R in a secondary E&R after 30 generations, and only confirmed the presence of selected alleles (Burny et al. 2020).

Our results suggest that large population sizes become especially crucial for short-term secondary E&R. The low power of short-term E&R with reduced experimental duration (Kofler & Schlötterer 2014) can be improved by a larger experimental population size. Our analyses show that to achieve satisfactory power after only 20 generations requires population sizes of tens of thousands of individuals– dimensions that are rather rare in current EE designs with strictly sexually reproducing organisms, where population sizes are mostly limited by the capacities to conduct such large experiments. Similar to previous simulation studies (Kofler & Schlötterer 2014; Kessner & Novembre 2015), we demonstrated that increasing the number of replicates results in more powerful secondary E&R. If large population sizes are not feasible, maintaining many replicates at small population size can help to boost secondary E&R performance. Additionally, the loss of one replicate in a highly replicated setup has a smaller impact than in an experimental setup with very few replicates. However, we show that additional replication cannot completely compensate the advantage of large population sizes in secondary E&R. Based on this observation, we conclude that independent of the crossing scheme, the power of secondary E&R will benefit from larger population sizes. Furthermore, the resolution will significantly increase with more replicates – a result that we anticipate can be generalized to different underlying genetic architectures as well as (model) organisms.

### Allele frequency estimation errors and read depth

The results of this study are based on true allele frequencies estimated without error - a rather optimistic assumption that will not hold for empirical data. In empirical E&R studies, numerous factors influence the accuracy of allele frequency estimates, such as the rate of sequencing errors, genome-wide read depth heterogeneity, and the average read depth (Baldwin-Brown et al. 2014; Kofler & Schlötterer 2014; Tilk et al. 2019).

Individual whole-genome sequencing for entire populations becomes quickly prohibitive with increasing sample size. Sequencing pools of individuals (Pool-Seq) (Schlötterer et al. 2014) provides a cost-effective approach that allows to robustly estimate allele frequency estimates and has become the method of choice for most E&R studies (Turner et al. 2011; Schlötterer et al. 2015). Since typical Pool-Seq studies combine all individuals from a given generation to reduce the sampling error (Schlötterer et al. 2014), the read depth of Pool-Seq studies will affect all experimental designs to the same extent as long as the coverage is considerably lower than the pool size.

### Choice of model organism

Our simulations are parameterized for *D. simulans* because this species is better suited for E&R studies than *D. melanogaster* (Barghi et al. 2017). While other species, such as *S. cerevisiae* (Burke et al. 2014) or *C. remanei* (Castillo et al. 2015) have been used to study adaptive response from standing genetic variation in outcrossing species, *Drosophila* is currently the most popular organism. Nevertheless, we anticipate that the results of this study can be generalized to other species, but the availability of sequenced inbred founder lines is probably a more severe restriction, which could limit the widespread use of secondary E&R.

### Selection regime

Our simulations are targeted for selection regimes with a moderate number of selection targets, with rather strong effects, as seen in empirical E&R studies using *Drosophila* (e.g. Mallard et al. 2018; Barghi et al. 2019; Michalak et al. 2019). Since strongly selected haplotype blocks typically have low starting frequencies, we consider the 1:1 crossing scheme a very realistic case, as most of the selection targets will be found only in a few ancestral founder lines. As outlined above, evolved haplotypes in dil:mt on the other hand will most likely carry multiple beneficial alleles that have recombined over the course of the primary E&R, which makes it challenging to fine-map one distinct selection target of interest as our results suggest.

Furthermore, we assumed the rather simple selection scenario of directional selection. More complex scenarios that include pleiotropy and/or epistasis were not considered. While both are very important factors that could affect the allele frequency changes in an E&R study, we would like to point out that secondary E&R is designed to study loci, which experienced a substantial allele frequency increase in a primary experiment, thus pleiotropic constraints are not expected to be a major confounding factor. Epistasis, on the other hand may be more pronounced in the 1:1 crossing scheme. It may be possible that the focal allele is only selected in a subset of the replicates, but not in others-or the response may differ between replicates. While this reduces the power of secondary E&R to fine-map the selection target, such a result may open the possibility to study the impact of epistatic effects. Since the founder lines will be available, it will be possible to repeat the experiment with replication for each genotype combination to assure that the heterogeneous response comes from epistatic effects and is not linked to stochastic changes.

Finally, it is important to keep in mind that the secondary E&R designs discussed here are not tailored to study highly polygenic traits, because in this study we were assuming that recombination facilitates the identification of a selection target with a major effect. For a highly polygenic trait, selected haplotype blocks identified in the primary E&R experiment (with multiple selected loci) would be broken down during the secondary E&R, which reduces their selective advantage. To what extent this different behavior in secondary E&R experiments can be used to distinguish these different architectures requires further work.

## Acknowledgements

We thank the members of the Institute of Population Genetics for fruitful discussion and support. Thanks to Roger Mundry who shared R scripts for the statistical analyses. Special thanks to Rupert Mazzucco, Robert Kofler, Kathrin Otte, and Christos Vlachos for their support regarding the computer simulations. Special thanks to Claire Burny and Robert Kofler for helpful comments on earlier versions of the manuscript. Last, but not least we are thankful to the anonymous reviewers for their constructive feedback. This work was supported by the Austrian Science Funds (FWF) [grant numbers W1225, P-32935]; and by the European Research Council (ERC) [grant ArchAdapt].

## Authors Contribution

AML, MD, and CS designed the analysis. AML performed the forward simulations, and the bioinformatic analysis. AML, and MD performed the statistical analysis. AML, MD, and CS wrote the manuscript.

## Supplementary Figures

**Figure S1.** Simulated versions of the 1:1 crossing scheme.

**Figure S2.** Number of ties in the 1:1_1f and dil:mt crossing schemes.

**Figure S3.** Power of the 1:1_1f and dil:mt crossing schemes after 20 generations of adaptation.

**Figure S4.** Resolution of the 1:1_1f and dil:mt crossing schemes after 20 generations of adaptation.

**Figure S5.** Allele frequency trajectories of selected SNPs in 1:1_2f.

**Figure S6.** Allele frequency trajectories of focal SNPs in 5 different crossing schemes.

**Figure S7.** Number of ties in 5 different crossing schemes.

**Figure S8.** Power of 5 different crossing schemes with 30 replicates per simulation.

**Figure S9.** Resolution of 5 different crossing schemes with 30 replicates per simulation.

## Supplementary Tables

**Table S1.** Number of SNPs per crossing scheme.

**Table S2.** Type II ANOVA (Model 1).

**Table S3.** Nagelkerke’s R^2^-index (Model 1).

**Table S4.** Type II ANOVA (Model 2).

**Table S5.** Nagelkerke’s R^2^-index (Model 2).

**Table S6.** Type II ANOVA (Model 3).

**Table S7.** Nagelkerke’s R^2^-index (Model 3).

---

**Figure S1.**
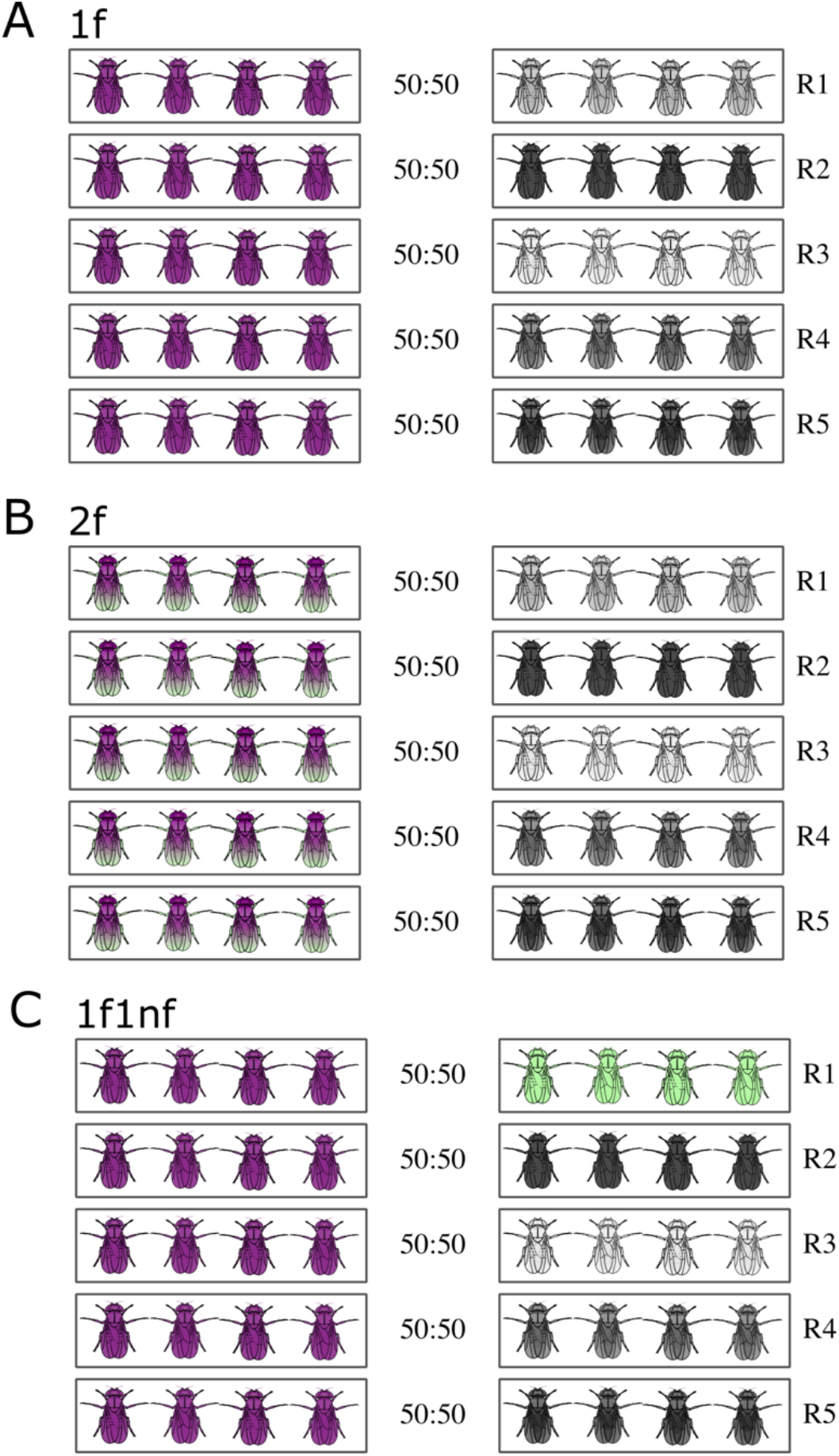
Simulated versions of the 1:1 crossing scheme. (A) Default version of the 1:1 crossing scheme (=**1f** for 1 focal SNP). Inbred flies with one target of selection (purple) are crossed to inbred flies without known beneficial variants. The starting frequency of each genotype is 50%. In each replicate the line with the beneficial allele (focal line, purple) is crossed to a different line lacking beneficial mutations (non-focal lines are colored in different shades of grey). Figure S1A is equal to Figure 1B in the main manuscript. (B) 2 focal SNPs (=**2f**) version of the 1:1 crossing scheme. The focal line carries two beneficial variants: the focal SNP we aim to fine map (purple) and one additional target of selection (green). (C) 1 focal, 1 non-focal SNP (=**1f1nf**) version of the 1:1 crossing scheme: One non-focal line carries on additional beneficial variant (green).

**Figure S2.**
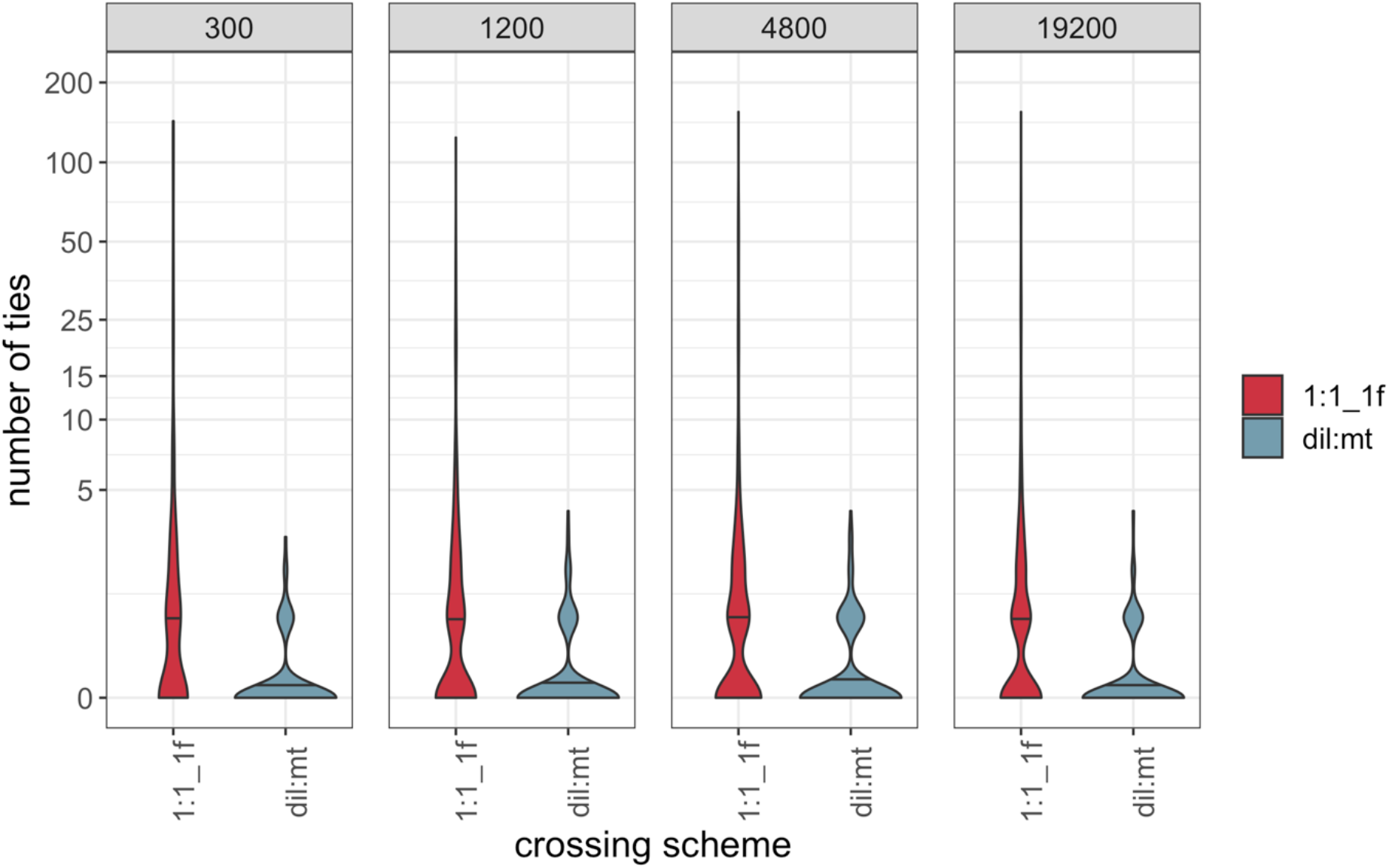
Violin plots of the number of ties (y axis = log_10_ (number of ties + 1)) for simulations where the true target of selection has the highest Cochran-Mantel-Haenszel (CMH) test statistic (success-A). The black horizontal lines in the violin plots display the median number of observed ties. Each panel shows the results for one particular simulated population size. Ties are defined as neighboring SNPs that have the same CMH test statistic as the target of selection.

**Figure S3.**
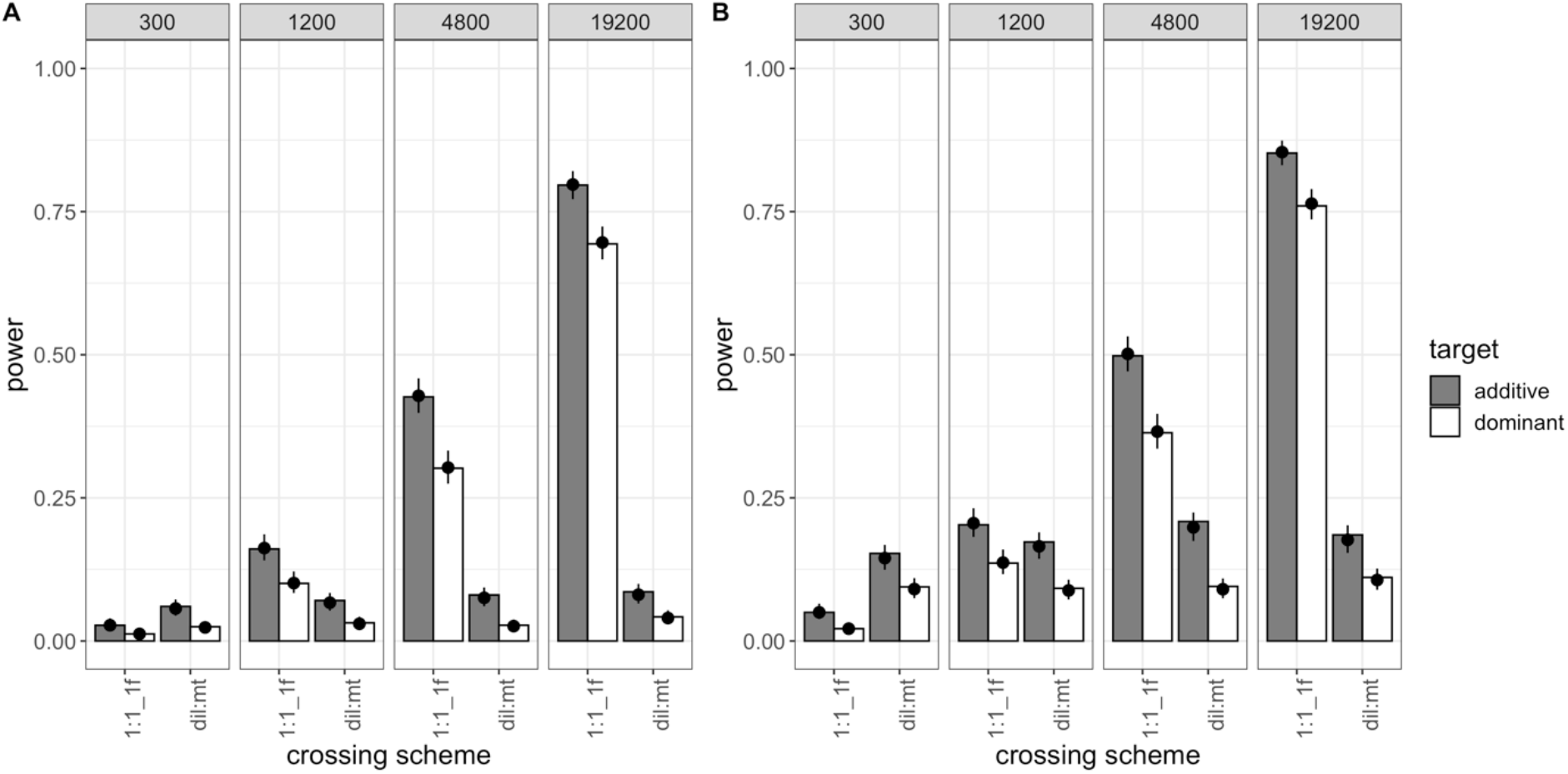
Power of the 1:1_1f and dil:mt crossing scheme at different population sizes (2,000 simulations/experimental design) after 20 generations of adaptation. Bars show the power (i.e., proportion of successful simulations) separately for each combination of crossing scheme (1:1_1f, dil:mt), population size (300; 1,200; 4,800; 19,200 individuals), and dominance coefficient (additive in grey, dominant in white). The dots with error bars display the model fit (Model 1) and its 95 % confidence interval. For the model fit, the selection coefficient was fixed to its global average, and combination-specific average starting allele frequencies were used. (A) shows the results for success-A (= selection target is the SNP with the highest Cochran-Mantel-Haenszel (CMH) test statistic) (B) shows the results for success-B (= selection target is not more than 100 SNPs away from the SNP with the highest CMH test statistic).

**Figure S4.**
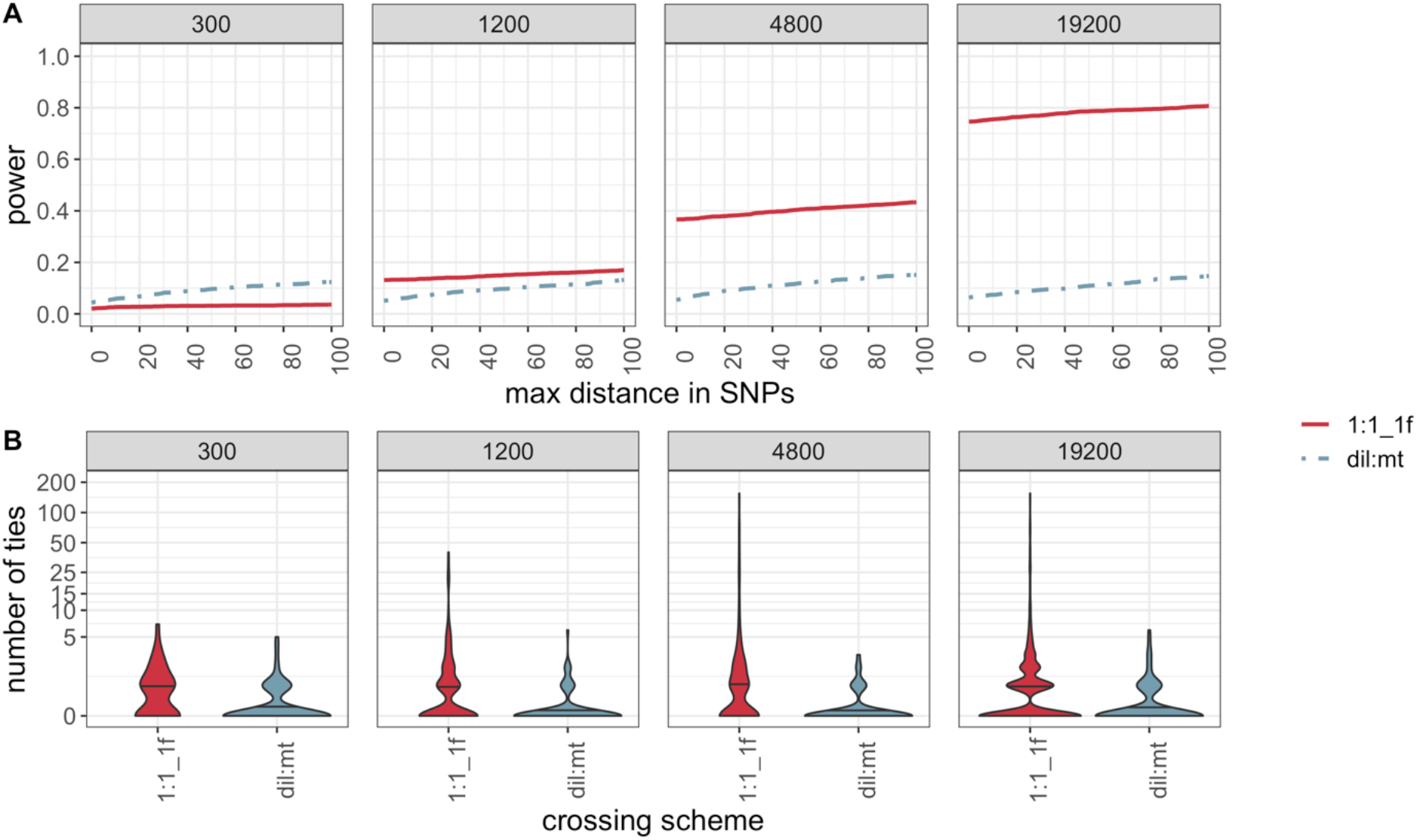
Resolution for the 1:1_1f and dil:mt crossing scheme simulated with different population sizes after 20 generations of adaptation. (A) Proportion of simulations (y-axis) that do not exceed a maximum distance in SNPs (x-axis) between the SNP with the highest Cochran-Mantel-Haenszel (CMH) test statistic and the true target of selection (B) Violin plots of the number of ties (y axis = log10 (number of ties + 1)) for simulations where the true target of selection has the highest CMH test statistic (success-A). The black horizontal lines in the violin plots display the median number of observed ties. Ties are defined as neighboring SNPs that have the same CMH test statistic as the target of selection.

**Figure S5.**
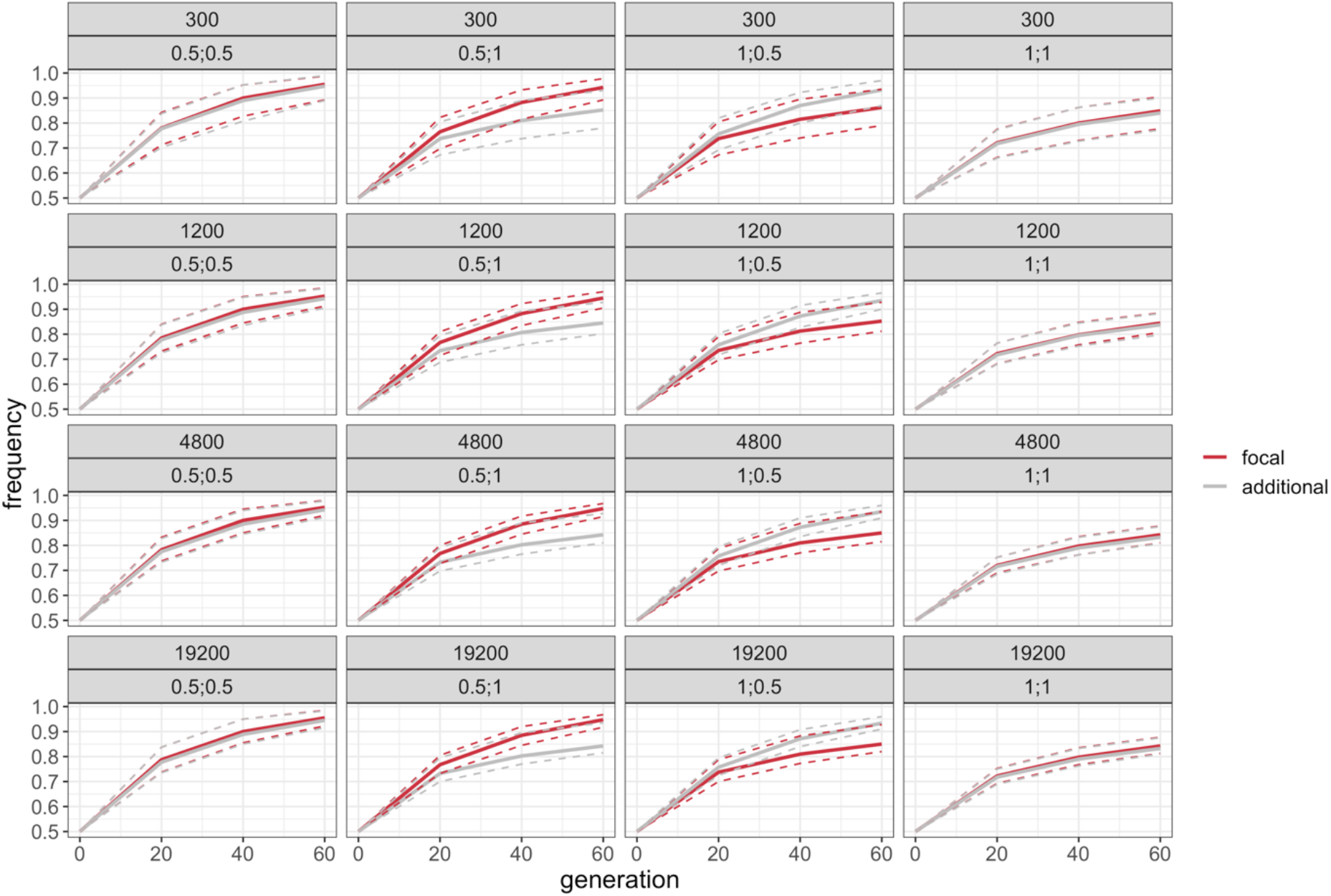
Median allele frequency trajectory of focal (red) and additional targets (grey) in 1:1_2f. For each single simulation, the target frequencies of a distinct generation were averaged over five replicates. Solid lines show the median allele frequency trajectory over 2,000 independent simulations per experimental design. Dashed lines show the 5 and 95 percentiles of the allele frequency trajectories. Each panel shows the median allele frequency trajectory of one particular dominance coefficient (focal target; additional target) and population size (300; 1,200; 4,800; 19,200 individuals) combination.

**Figure S6.**
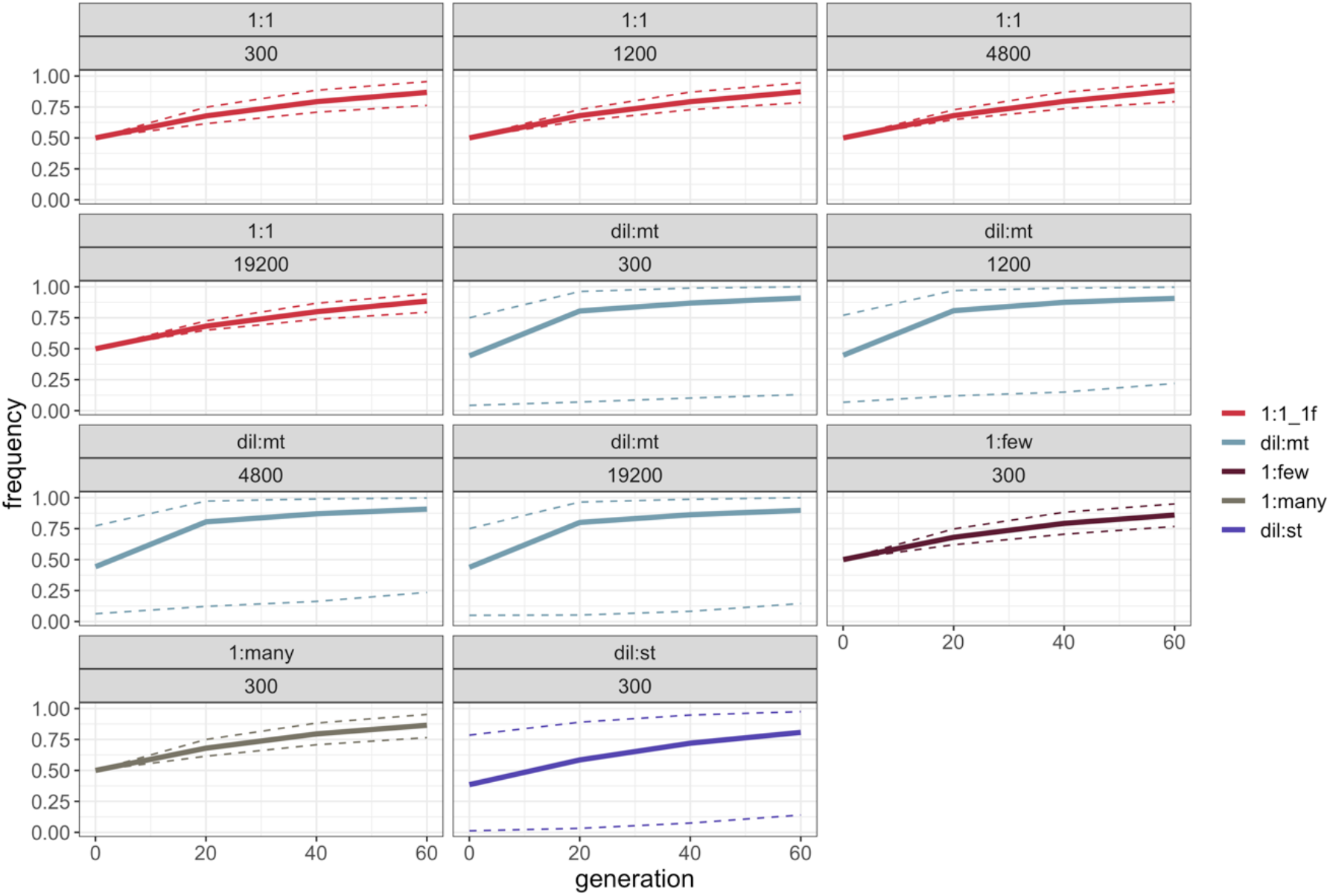
Median allele frequency trajectory of focal targets in different experimental designs. For each single simulation, the focal target frequency of a distinct generation was averaged over five replicates. Solid lines show the median allele frequency trajectory over 2,000 independent simulations per experimental design. Dashed lines show the 5 and 95 percentiles of the allele frequency trajectories. Each panel shows the median allele frequency trajectory of one particular crossing scheme (1:1_1f, dil:mt, 1:few, 1:many, dil:st) and population size (300; 1,200; 4,800; 19,200 individuals) combination.

**Figure S7.**
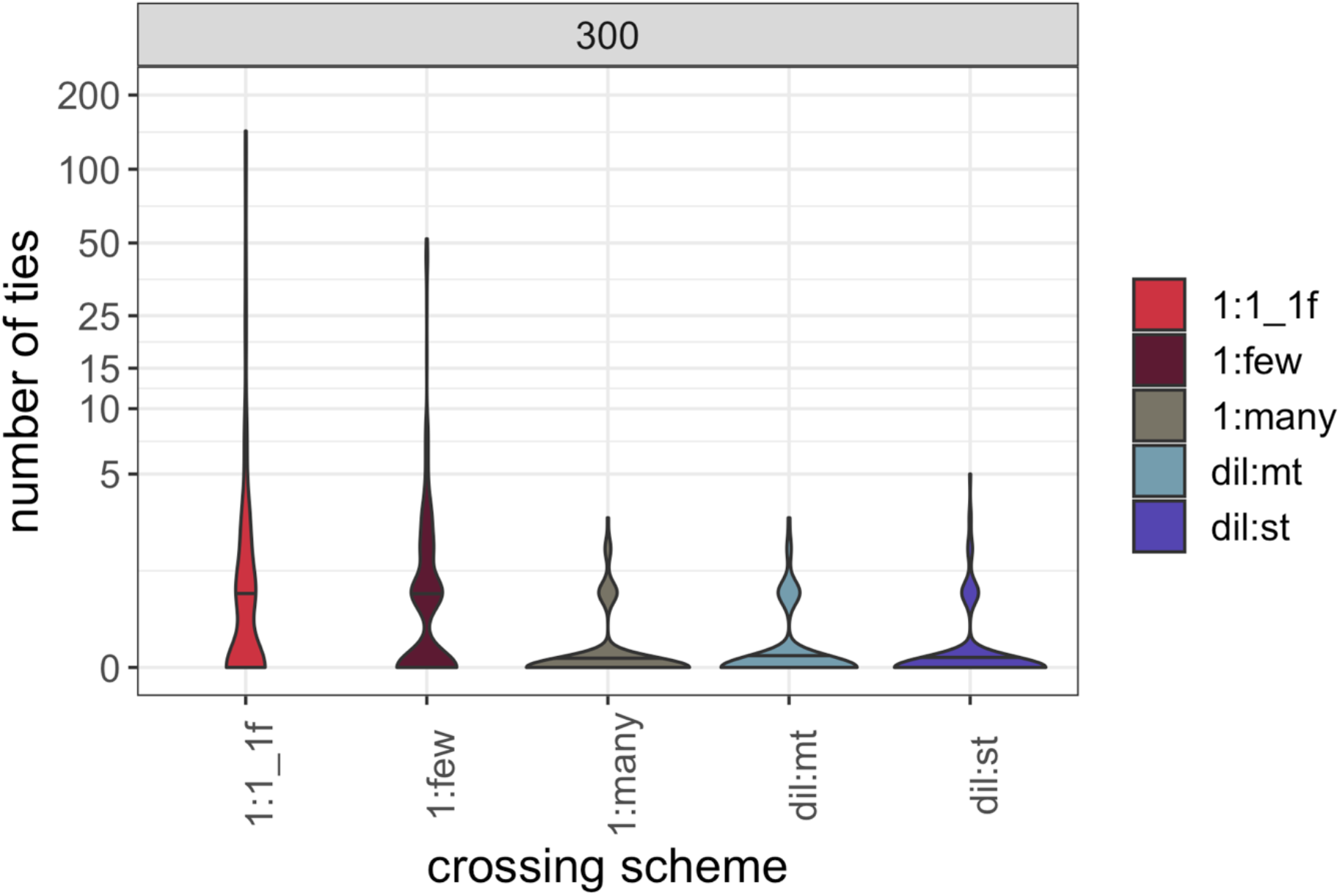
Violin plots of the number of ties (y axis = log10 (number of ties + 1)) for simulations where the true target of selection has the highest Cochran-Mantel-Haenszel (CMH) test statistic (success-A, population size = 300 individuals; 5 replicates; 2,000 simulations/experimental design). The black horizontal lines in the violin plots display the median number of observed ties. Ties are defined as neighboring SNPs that have the same CMH test statistic as the target of selection.

**Figure S8.**
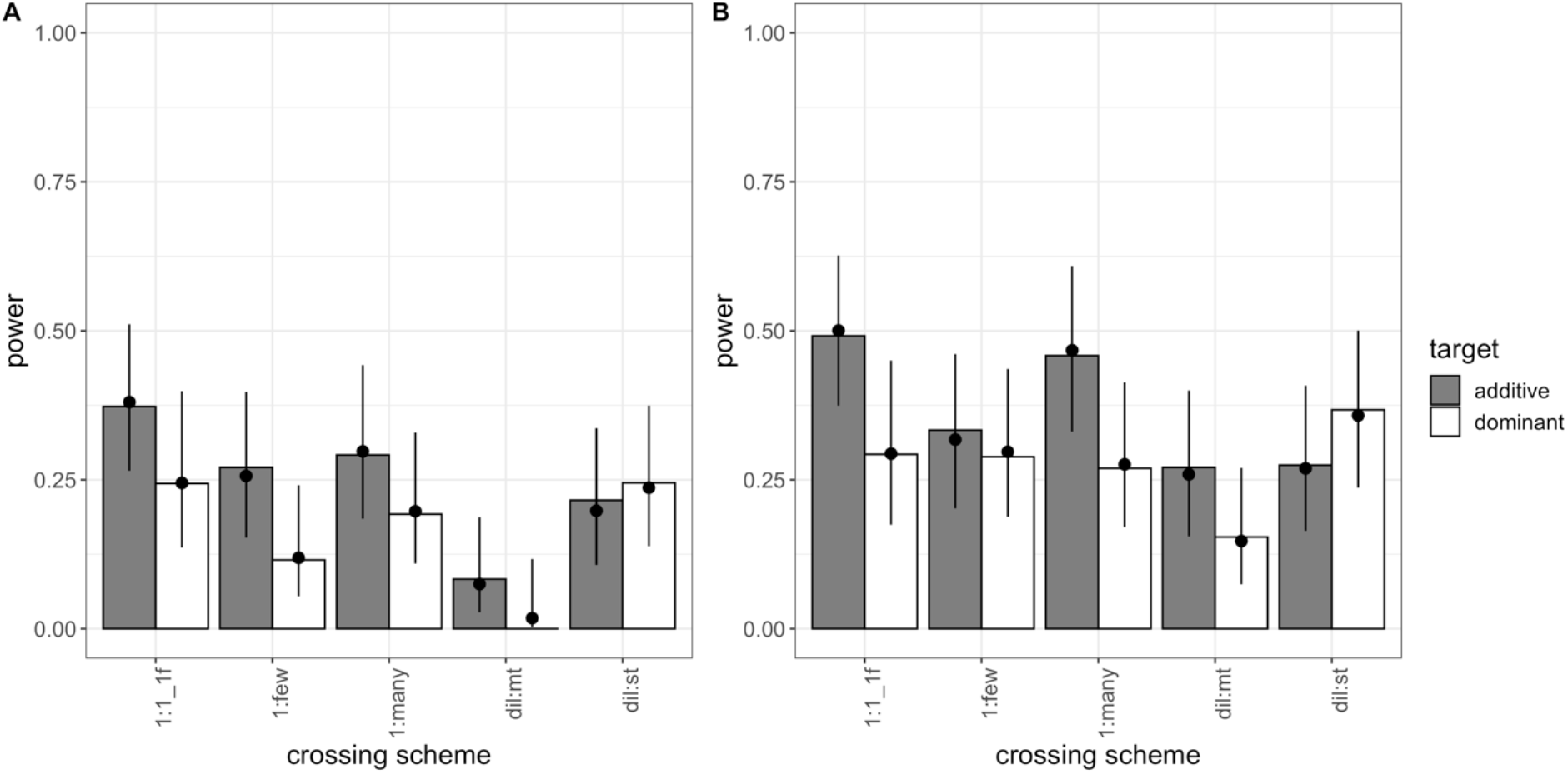
Power of five different crossing schemes (population size = 300 individuals; 30 replicates; 100 simulations/experimental design). Bars show the power (i.e., proportion of successful simulations) separately for each combination of crossing scheme and dominance coefficient (additive in grey; dominant in white). The dots with error bars display the estimate from the fitted model (Model 3) and its 95 % confidence interval. For the model fit, the selection coefficient was fixed to its global average, and combination-specific average starting allele frequencies were used. (A) shows the results for success-A (= selection target is the SNP with the highest Cochran-mantel-Haenszel (CMH) test statistic), (B) shows the results for success-B (= selection target is not more than 100 SNPs away from the SNP with the highest CMH test statistic).

**Figure S9.**
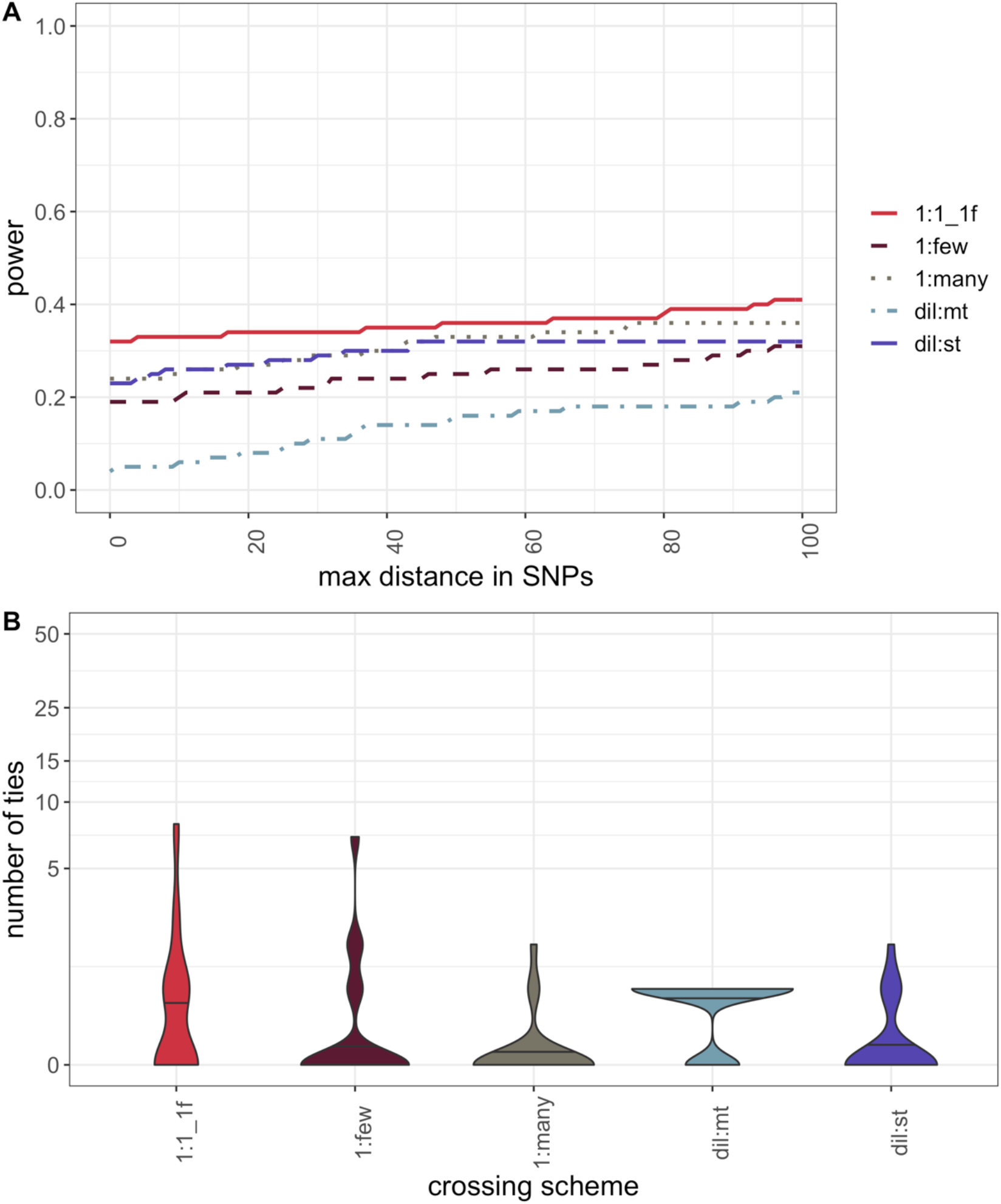
Resolution for five different crossing schemes (population size = 300 individuals; 30 replicates; 100 simulations/experimental design). (A) Proportion of simulations (y-axis) that do not exceed a maximum distance in SNPs (x-axis) between the SNP with the highest Cochran-Mantel-Haenszel (CMH) test statistic and the true target of selection (B) Violin plots of the number of ties (y axis = log_10_ (number of ties + 1)) for simulations where the true target of selection has the highest CMH test statistic (success-A). The black horizontal lines in the violin plots display the median number of observed ties. Ties are defined as neighboring SNPs that have the same CMH test statistic as the target of selection.

**Table S1.** Number of SNPs per crossing scheme on chromosome-arm 2L.

| **crossing scheme** | **median** | **range** |
| --- | --- | --- |
| 1:1 | 523,008 | 479,507 – 527,771 |
| 1:few | 526,510 | 522,470 – 530,230 |
| 1:many | 970,466 | 970,466 – 970,466 |
| dil:mt | 970,466 | 970,466 – 970,466 |
| dil:st | 970,466 | 970,466 – 970,466 |

**Table S2.**
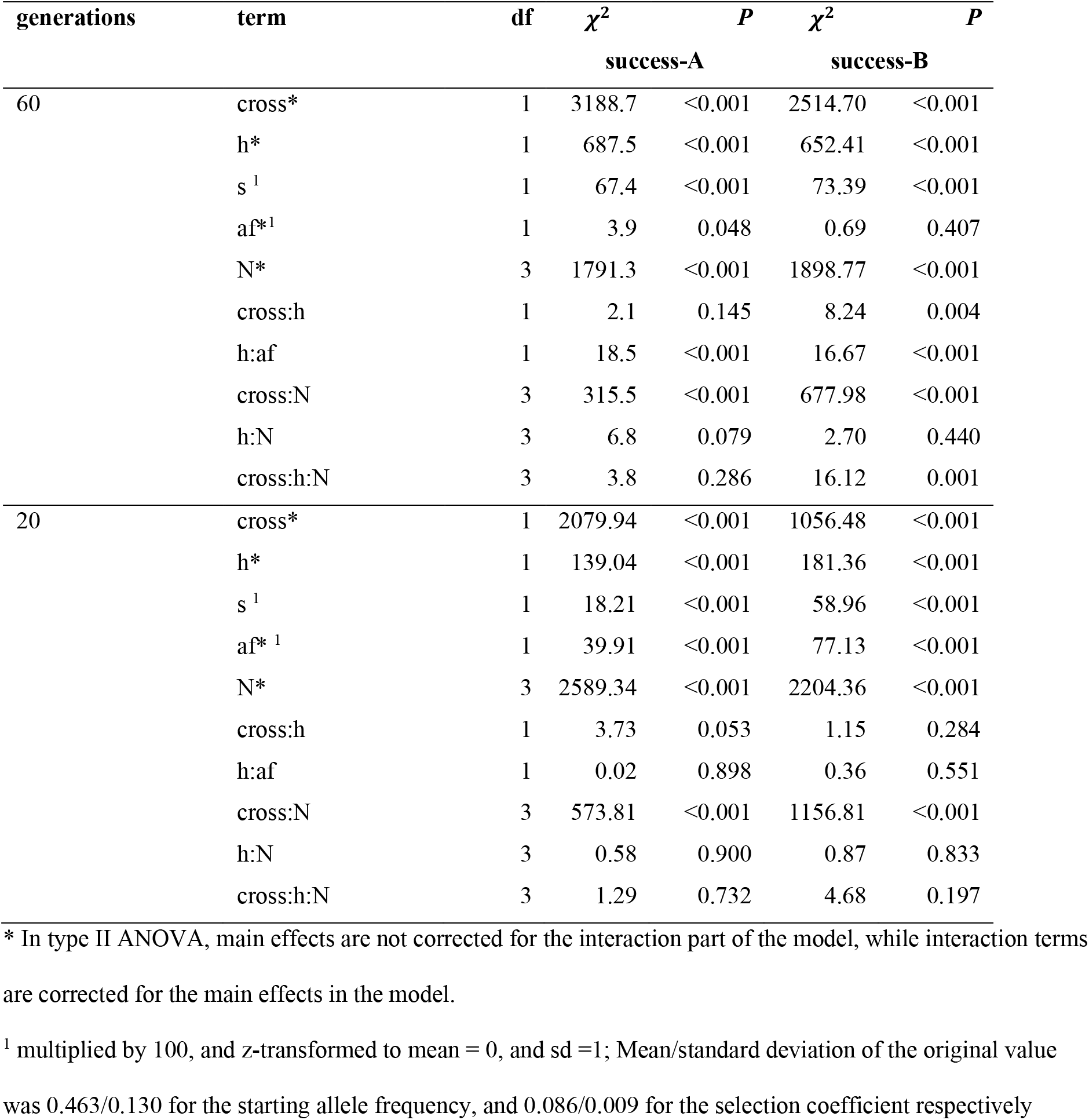
Type II ANOVA of explanatory variables for the analysis of success in secondary Evolve and Resequence studies with model 1 after 60 and 20 generations of adaptation. cross= crossing scheme; h= dominance coefficient; s= selection coefficient; af= mean starting allele frequency of the focal target over 5 replicates; N= population size; cross:h = interaction term between crossing scheme and dominance coefficient; h:af = interaction term between dominance coefficient and mean starting allele frequency; cross:N = interaction term between crossing scheme and population size; h:N = interaction term between dominance coefficient and population size; cross:h:N = interaction term between crossing scheme, dominance coefficient, and population size.

**Table S3.** Nagelkerke’s R^2^-index of each explanatory variable including all its interactions in the logistic regression (Model 1). If the reduced model explains the data as well as a full model that includes the evaluated effect, Nagelkerke’s R^2^-index is 0.

| **generations** | **effect** | **R<sup>2</sup>**<br>**success-A** | **R<sup>2</sup>**<br>**success-B** |
| --- | --- | --- | --- |
| 60 | crossing scheme | 0.305 | 0.264 |
|  | dominance coefficient | 0.077 | 0.069 |
|  | selection coefficient | 0.007 | 0.007 |
|  | mean starting allele frequency | 0.003 | 0.002 |
|  | population size | 0.204 | 0.224 |
| 20 | crossing scheme | 0.283 | 0.211 |
|  | dominance coefficient | 0.019 | 0.021 |
|  | selection coefficient | 0.002 | 0.007 |
|  | mean starting allele frequency | 0.005 | 0.009 |
|  | population size | 0.322 | 0.295 |

**Table S4.** Type II ANOVA of explanatory variables for the analysis of success in secondary Evolve and Resequence studies with model 2. architecture= version of the 1:1 crossing scheme in combination with the dominance coefficient of the additional target; h= dominance coefficient of the selection target of interest; s= selection coefficient; N= population size; architecture:h = interaction term between architecture and dominance coefficient of the selection target of interest; architecture:N = interaction term between architecture and population size; h:N = interaction term between dominance coefficient of the selection target of interest and population size; architecture:h:N = interaction term between architecture, dominance coefficient of the selection target of interest, and population size.

| term | df | $\chi^2$ | P | $\chi^2$ | P |
| --- | --- | --- | --- | --- | --- |
|  |  | success-A |  | success-B |  |
| architecture* | 4 | 855.4 | <0.001 | 1509.9 | <0.001 |
| h* | 1 | 2742.4 | <0.001 | 2755.9 | <0.001 |
| s<sup>1</sup> | 1 | 181.9 | <0.001 | 137.5 | <0.001 |
| N* | 3 | 5230.6 | <0.001 | 6661.9 | <0.001 |
| architecture:h | 4 | 697.8 | <0.001 | 708.6 | <0.001 |
| architecture:N | 12 | 17.5 | 0.132 | 42.7 | <0.001 |
| h:N | 3 | 12.3 | 0.006 | 3.0 | 0.399 |
| architecture:h:N | 12 | 27.9 | 0.006 | 29.8 | 0.003 |

\* In type II ANOVA, main effects are not corrected for the interaction part of the model, while interaction terms are corrected for the main effects in the model.
<sup>1</sup> multiplied by 100, and z-transformed to mean = 0, and sd =1; Mean/standard deviation of the original value was 0.085/0.009.

**Table S5.**
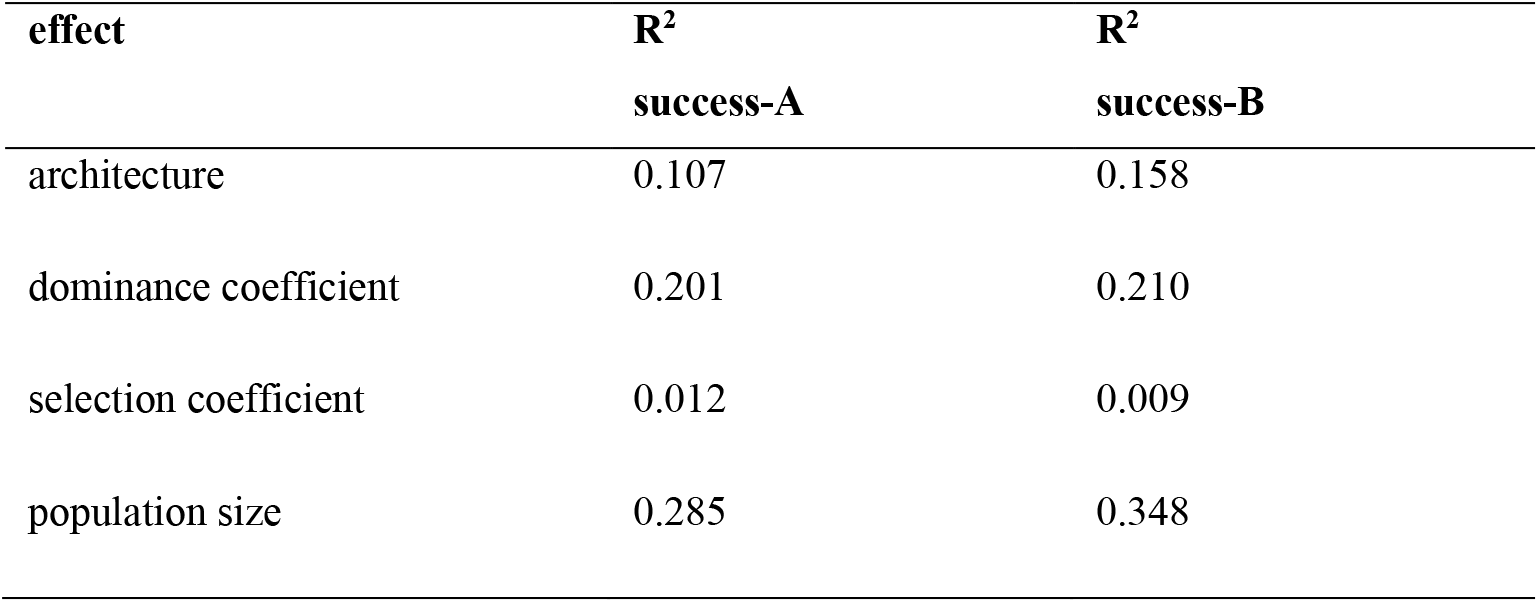
Nagelkerke’s R^2^-index of each explanatory variable including all its interactions in the logistic regression (Model 2). If the reduced model explains the data as well as a full model that includes the evaluated effect, Nagelkerke’s R^2^-index is 0.

| **effect** | **$R^2$**<br>**success-A** | **$R^2$**<br>**success-B** |
| --- | --- | --- |
| architecture | 0.107 | 0.158 |
| dominance coefficient | 0.201 | 0.210 |
| selection coefficient | 0.012 | 0.009 |
| population size | 0.285 | 0.348 |

**Table S6.** Type II ANOVA of explanatory variables for the analysis of success in secondary Evolve and Resequence studies with model 3. cross= crossing scheme; h= dominance coefficient; s= selection coefficient; af= mean starting allele frequency of the focal target over 5 replicates; cross:h= interaction term between crossing scheme and dominance coefficient; h:af= interaction term between dominance coefficient and mean starting allele frequency.

| generations | term | df | $\chi^2$ | P | $\chi^2$ | P |
| --- | --- | --- | --- | --- | --- | --- |
|  |  |  | success-A |  | success-B |  |
| 5 | cross* | 4 | 218.65 | <0.001 | 93.01 | <0.001 |
|  | h* | 1 | 371.35 | <0.001 | 321.13 | <0.001 |
|  | s<sup>1</sup> | 1 | 76.20 | <0.001 | 72.09 | <0.001 |
|  | af*<sup>1</sup> | 1 | 5.55 | 0.019 | 12.66 | <0.001 |
|  | cross:h | 4 | 77.19 | <0.001 | 97.21 | <0.001 |
|  | h:af | 1 | 7.56 | 0.006 | 7.07 | 0.008 |
| 30 | cross* | 4 | 28.79 | <0.001 | 11.28 | 0.024 |
|  | h* | 1 | 5.13 | 0.024 | 4.40 | 0.036 |
|  | s<sup>1</sup> | 1 | 2.63 | 0.105 | 3.65 | 0.056 |
|  | af*<sup>1</sup> | 1 | 1.34 | 0.247 | 0.13 | 0.716 |
|  | cross:h | 4 | 2.68 | 0.612 | 5.41 | 0.247 |
|  | h:af | 1 | 1.73 | 0.189 | 1.01 | 0.315 |

\* In type II ANOVA, main effects are not corrected for the interaction part of the model, while interaction terms are corrected for the main effects in the model.
<sup>1</sup> multiplied by 100, and z-transformed to mean = 0, and sd =1; Mean/standard deviation of the original value were 0.463/0.133 for the starting allele frequency, and 0.085/0.009 for the selection coefficient respectively.

**Table S7.**
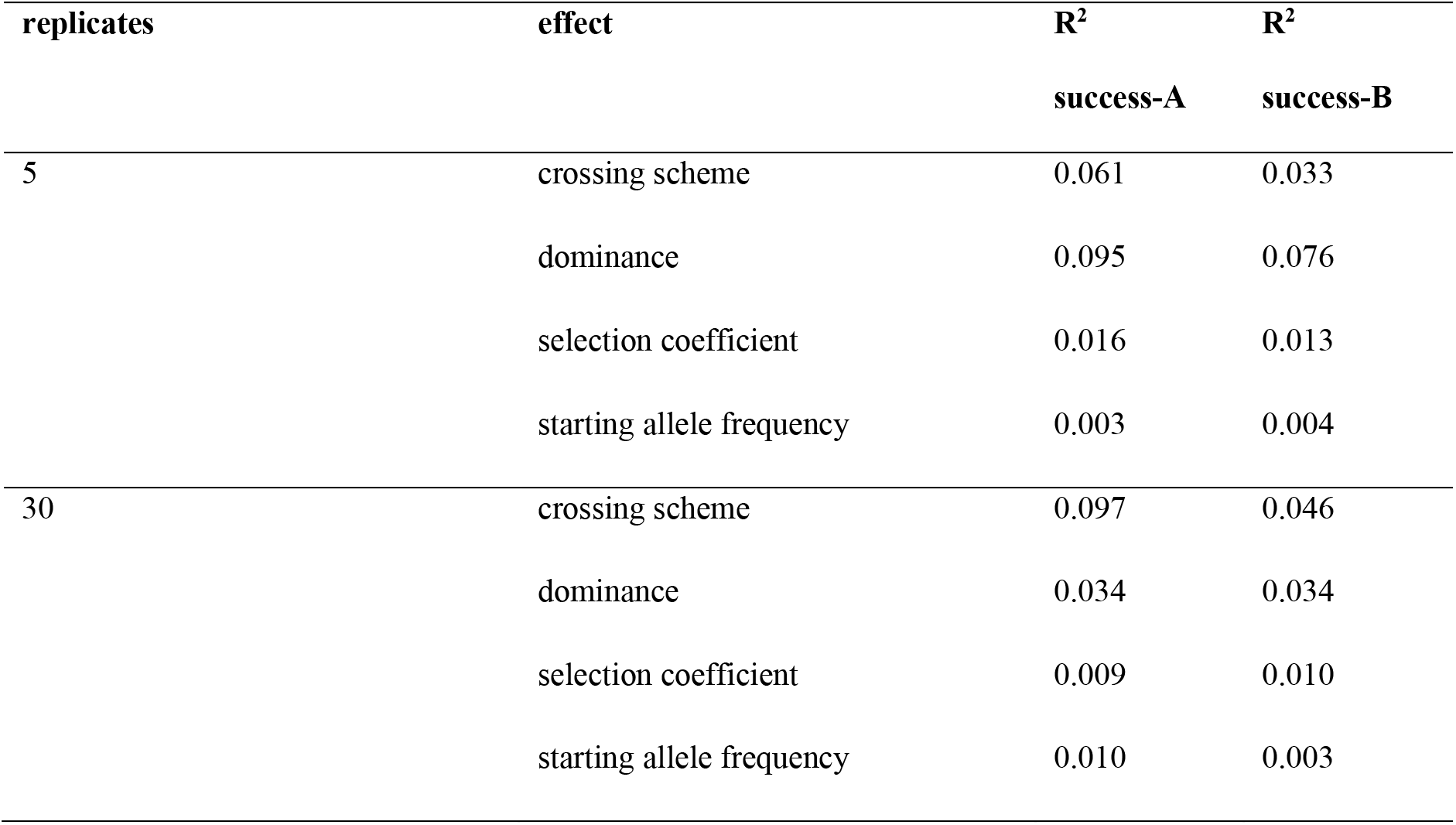
Nagelkerke’s R^2^-index of each explanatory variable including all its interactions in the logistic regression (Model 3). If the reduced model explains the data equally well as a full model that includes the evaluated effect, Nagelkerke’s R^2^-index is 0.

| replicates | effect | $R^2$ | $R^2$ |
| --- | --- | --- | --- |
|  |  | success-A | success-B |
| 5 | crossing scheme | 0.061 | 0.033 |
|  | dominance | 0.095 | 0.076 |
|  | selection coefficient | 0.016 | 0.013 |
|  | starting allele frequency | 0.003 | 0.004 |
| 30 | crossing scheme | 0.097 | 0.046 |
|  | dominance | 0.034 | 0.034 |
|  | selection coefficient | 0.009 | 0.010 |
|  | starting allele frequency | 0.010 | 0.003 |

